# A mechanohydraulic model supports a role for plasmodesmata in cotton fiber elongation

**DOI:** 10.1101/2023.06.30.547211

**Authors:** Valeria Hernández-Hernández, Olivier C. Marchand, Annamaria Kiss, Arezki Boudaoud

## Abstract

Plant cell growth depends on turgor pressure, the cell hydrodynamic pressure, which drives expansion of the extracellular matrix (the cell wall). Turgor pressure regulation depends on several physical, chemical and biological factors, including: vacuolar invertases, which modulate osmotic pressure of the cell, aquaporins, which determine the permeability of the plasma membrane to water, cell wall remodeling factors, which determine cell wall extensibility (inverse of effective viscosity), and plasmodesmata, which are membrane-lined channels that allow free movement of water and solutes between cytoplasms of neighbouring cells, like gap junctions in animals. Plasmodesmata permeability varies during plant development and experimental studies have correlated changes in the permeability of plasmodesmal channels to turgor pressure variations. Here we study the role of plasmodesmal permeability in cotton fiber growth, a type of cell that increases in length by at least 3 orders of magnitude in a few weeks. We incorporated plasmodesma-dependent movement of water and solutes into a classical model of plant cell expansion. We performed a sensitivity analysis to changes in values of model parameters and found that plasmodesmal permeability is among the most important factors for building up turgor pressure and expanding cotton fibers. Moreover, we found that non-monotonic behaviors of turgor pressure that have been reported previously in cotton fibers cannot be recovered without accounting for dynamic changes of the parameters used in the model. Altogether, our results suggest an important role for plasmodesmal permeability in the regulation of turgor pressure.

**Significance Statement:** The cotton fiber is among the plant cells with the highest growth rates. In cultivars, a single fiber cell generally reaches a few centimeters in length. How such size is achieved is still poorly understood. In order to tackle this question, we built a comprehensive mathematical model of fiber elongation, considering cell mechanics and water entry into the cell. Model predictions agree with experimental observations, provided that we take into account active opening and closure of plasmodesmata, the nano-channels that connect the fiber with neighboring cells. Because cotton fiber length is a key factor for yarn quality, our work may help understanding the mechanisms behind an important agronomic trait.

---

**E**xpansion of the plant cell involves mechanical and hydraulic processes. Mechanical processes include the ability of the cell wall to increase in surface area, called wall extensibility. Hydraulic processes include water movement across the plasma membrane, through aquaporins, or between cells, through channels known as plasmodesmata, that create cytoplasmic continuity between cells, like gap junctions in animals. James Lockhart (1) developed a model that has become widely used in the study of the mechano-hydraulic processes behind irreversible plant expansion. In its original form, Lockhart did not account for plasmodesmata permeability. It has been shown that the diameter of the channel (therefore also its permeability) may vary during plant development which affects movement between cells of small molecules like sucrose (2). The idea that plasmodesmata can regulate fluxes of solutes and water has led to the hypothesis that plasmodesmal permeability may be important for building up turgor during cell expansion. As a consequence of this, theoretical studies have addressed the hydraulic conductivity and the permeability to solutes of a single plasmodesma (3–6) and have started to integrate the role of plasmodesmal permeability into models of plant cell expansion (7). Likewise, gap junction permeability was recently accounted for in models of volume regulation in animal cells (8). Here we further consider the role of plasmodesmal permeability in the expansion of plant cells.

The cotton fiber is an ideal system to study the regulation of cell expansion because they are single epidermal cells that mostly increase in length (9, 10). There are several species of cotton (*Gossypium*), which enables comparisons between phenotypes. In *Gossypium hirsutum*, fibers start growing on the day of anthesis until 20-26 days after anthesis (DAA) (9, 10). A study performed by Ruan *et al*. (11, 12) reported that plasmodesmata change permeability from open (0-9 DAA) to closed (10-15 DAA), and then to open (16 onwards) again during fiber growth. The timing of this pattern may depend on cultivars and cotton species (13). Besides the dynamics of plasmodesmal permeability, Ruan and colleagues (11) also reported peak values of turgor and osmotic pressures at around 15 DDA, which correlated with the closure of plasmodesmata. The authors also found that the turgor pressure difference between the cotton fiber cell and its adjacent seed coat cell is largest when plasmodesmata are closed. Interestingly, a recent multicellular model of equivalent cells shows that low cell-to-cell permeability increases the turgor pressure differences between neighboring cells (7), which motivated us to model cotton fiber elongation.

Several processes may contribute to an increase in cell osmotic pressure during fiber growth: an increase in the expression level of sucrose and K+ transporters (11), and of vacuole invertase 1 (VIN1) (14); an increase in potassium and malate concentrations (15). While VIN1 and solute transporters might enhance the accumulation of solutes within the fiber, closure of plasmodesmata may prevent leakage of solutes and of water causing a rise of turgor (11). In order to better understand the role of plasmodesmata, we study whether a minimal hydro-mechanical model can reproduce the observations of turgor and osmotic pressure peaks during fiber growth. Furthermore, we investigate whether the observed correlation of these peaks with plasmodesmata closure and increase in solute concentrations are causal in this model.

## Results and discussion

### Mechano-hydraulic model of the growing cotton fiber

We consider a single cotton fiber, connected to neighboring cells, as depicted in Figure 1. We aim at a model that is amenable to a comprehensive exploration of the parameter space. We thus approximate the fiber geometry to a cylinder of radius *r* and length *l*. Given that cotton fibers undergo a huge increase in length and moderate changes in diameter, we assume that the fiber grows only in length. In the following, we derive the system of differential equations that governs the dynamics of the volume of the growing cotton fiber cell, *V* = *πr*^2^*l*, of its osmotic pressure, *π*_fiber_, and of its mechanical (turgor) pressure, *P*_fiber_.

**Fig. 1.**
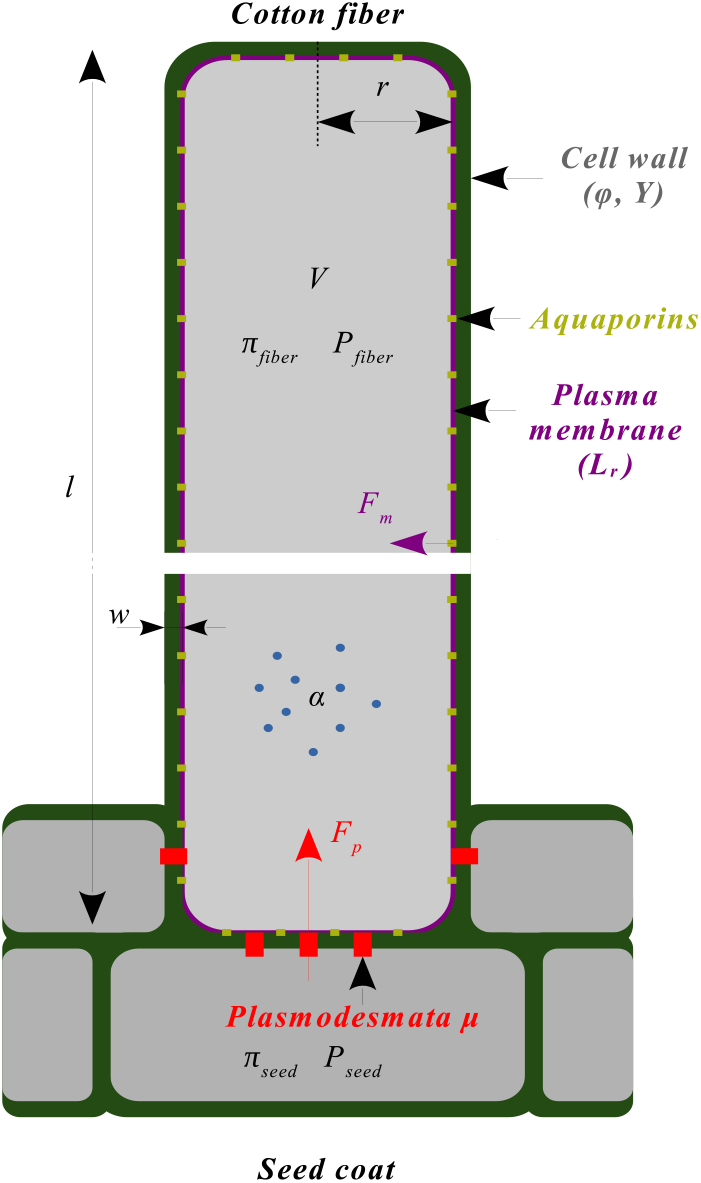
Mechano-hydraulic model of a cotton fiber. Cell shape is approximated by a cylinder of radius *r*, length *L*, and volume *V*. The cell wall has thickness *w*, extensibility *φ*, and yield stress *Y*. The cell is characterized by an osmotic pressure *π*_fiber_, a mechanical (turgor) pressure *P*_fiber_, and a rate of solute import/synthesis *α*. The plasma membrane has hydraulic conductivity *L*_*r*_, mainly associated with aquaporins. Water moves between the cell and the outside through two pathways: through the plasma membrane (apoplastic pathway) and through plasmodesmata, the nanometric channels connecting the fiber to neighboring cells (symplastic pathway). The three main variables that describe the fiber cell are its volume, *V*, its turgor pressure, *P*_fiber_, and its osmotic pressure *π*_fiber_.

### Water dynamics

Since water is nearly incompressible, the observed change in volume *V* of the cell is due to water moving in or out of the cell. Water can move either through the plasma membrane (mostly through aquaporins), with flux *F*_*m*_, or through plasmodesmata, with flux *F*_*p*_:

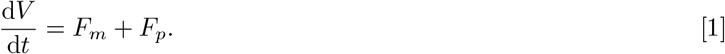

The flux through the membrane is driven by the difference in water potential (chemical potential of water) between the cell (Ψ_*fiber*_) and the external environment (Ψ_apoplasm_), ΔΨ = Ψ_*fiber*_ *−* Ψ_apoplasm_. Water flows towards the compartment with a lower potential

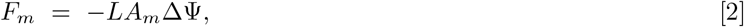

where *L* is the membrane permeability (per unit area) and *A*_*m*_ is the membrane surface. It is commonly held that water moves through and along all cell walls (16). In particular, we consider that water is available all around the fibers, which do not seem to have a cuticle (17), and that the water potential of the apoplasm is constant all along the fiber. We keep this assumption for the side walls of the cotton fiber, and we assume that water may arrive and cross the membrane on all sides of the cell. Because the fibers are very long, the area of cell ends is much smaller than the lateral area, and thus we take *A*_*m*_ = 2Π*rl* = 2*V /r*, which is proportional to the volume of the cell *V* .

Water potential in a compartment combines mechanical pressure (turgor, *P*), and osmotic pressure (*π*) in this compartment: Ψ = *P − π* (16). We here consider the developmental phase during which the fiber elongates, while the remainder of the seed has ceased expansion. Accordingly, neighboring cells are at thermodynamic equilibrium with the apoplasmic space, Ψ_seed_ = Ψ_apoplasm_, so that the water potential difference, ΔΨ = Δ*P −* Δ*π*, with Δ*P* = *P*_fiber_ *− P*_seed_ and Δ*π* = *π*_fiber_ *− π*_seed_. The water flux through the membrane is then

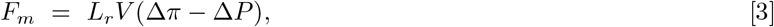

where *L*_*r*_ = 2*L/r* is the relative hydraulic conductivity of the membrane (1, 18, 19). An increase in fiber turgor pressure, *P*_fiber_, leads to a decrease of water flux, whereas an increase in fiber osmotic pressure, *π*_fiber_, leads to an increase of water flux towards the cell.

Water also flows through plasmodesmata channels linking the fiber cell to the seed cell, depending on the turgor pressure difference between the two compartments,

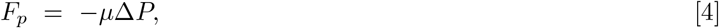

where *μ* is the total permeability associated with plasmodesmata.

Altogether, the rate of fiber volume change is given by Equations 1, 3 and 4, which require prescribing the dynamics of osmotic and turgor pressure.

### Solute dynamics

The dynamics of osmotic pressure depends on solute dynamics. The total number of solute particles *N* in the fiber changes through two processes. On the one hand, the fiber may exchange solutes with neighboring cells through plasmodesmata. The concentration of solutes transported by the flux (given by Eq. 4) depends on the flux direction: if the hydrostatic pressure in the seed is higher than in the fiber, the solutes in the seed, with concentration *c*_*seed*_, enter the fiber, otherwise the solutes in the fiber, with concentration *c*_*fiber*_, leave the fiber. On the other hand, the fiber cell uptakes solutes from the cell wall compartment or breaks solute particles into smaller ones (thanks for instance to invertases), leading to an increase in the number of solute particles. As a consequence, the rate of change of solute number is given by

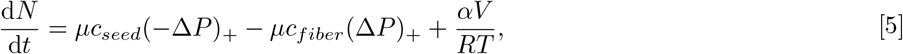

where (*x*)_+_ denotes the positive part of *x*, and we introduced the constant of perfect gases *R*, temperature *T*, and a normalized rate of solute increase *α*. Finally, we relate any solute concentration *c* to osmotic pressure *π* in the corresponding compartment by *π* = *NRT/V* = *cRT*. This approximation is valid for low concentrations, and we obtain in this case the following equation for the dynamics of the osmotic pressure in the fiber

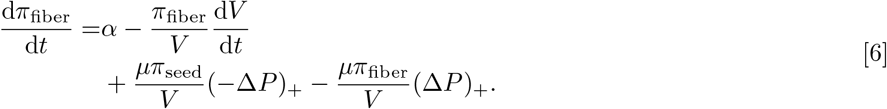

### Cell wall expansion

The cell wall is under tensile stress due to turgor pressure. This tension leads to both elastic (reversible) and plastic-like (irreversible) deformation of the wall. Given that typical values of longitudinal elastic deformation are smaller than 10%, even for soft cell walls (24), the huge increase in fiber cell length is mostly associated with irreversible deformation. Therefore we have chosen to neglect elastic deformations.

Cotton fibers grow diffusely (all along their length) at early stages (25, 26). We assume that this holds up to growth arrest and we use the classical Lockhart’s equation to model elongation. When turgor is above a critical yield threshold (*Y*), the rate of volume increase is determined by turgor in excess of *Y* and cell wall extensibility, *φ*,

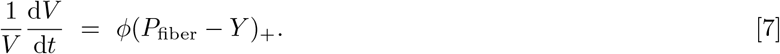

Here *Y* and *φ* are fixed values that are representative of parameters that potentially vary along the fiber. Equations 3 and 7 would form together the complete version of the Lockhart equation (1, 27, 28) in which growth is potentially limited by wall extensibility and by membrane conductivity. Here, in addition, we account for cytoplasmic flow through plasmodesmata (Equation 4). Using Eq. (1) that describes the total flux of water into the cell, we eliminate the turgor pressure *P*_fiber_ and we obtain the pressure difference Δ*P* = *P*_fiber_ *− P*_seed_ as

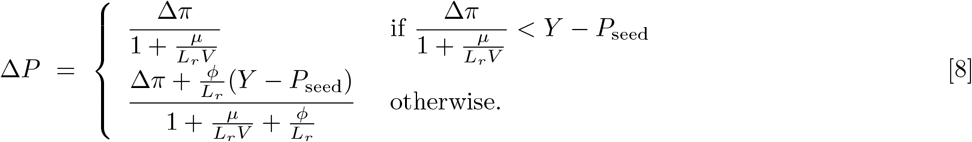

The first line of Equation 8 holds when there is no fiber growth – the inequality reduces to the classical condition of no growth when plasmodesmata are closed (*μ* = 0) – and the second line applies to when the fiber grows. Equations 1 and 6, together with Eqs. 3, 4, and 8 form a system of differential equations for the two variables *V* and *π*_fiber_, with seven parameters: plasma membrane permeability *L*_*r*_, hydraulic conductivty of all plasmodesmataμ, solute uptake/synthesis rate *α*, extensibility *φ*, seed turgor pressure *P*_seed_, seed osmotic pressure *π*_seed_, and yield threshold *Y*. We consider the initial conditions *V* (*t* = 0) = *V* (0) and *π*_fiber_(*t* = 0) = *π*_seed_.

### Model predictions for constant parameters

#### Reference values

We start with the analysis of the model with constant parameters, and in particular, we choose as reference parameter values the geometric mean of the range limits given in Table 1. We consider these ranges to be the biologically relevant values of parameters. As indicated in Table 1, these values were either found directly in the literature or were estimated from the literature. The methodology of estimation is detailed in the Supplementary note. For instance, the estimation of plasmodesmal permeabilityμ is based on the methodology used in (5), and accounts for a range of possible geometries and dimensions of plasmodesmata channels. In the following, we refer to the model with constant parameters taking these reference values as the reference model. The volume, the osmotic and turgor pressures, and fluxes of the reference model with the initial volume *V* (0) = 1.88 *×* 10*−*4 mm^3^ are plotted in figure 2. For these reference parameters, we see that both osmotic and turgor pressures reach constant values (after about 400 h) after a monotonic regime. Only the behavior of the first few hours depends on the initial conditions. The growth rate defined as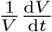 becomes constant as soon as the pressures reach their limiting values.

**Table 1.** Model parameters.

| Symbol | Meaning and unit | Range | Reference value | Relative deviation | Source(s) |
| --- | --- | --- | --- | --- | --- |
| $L_r$ | Normalized hydraulic conductivity of the cell membrane ($Pa^{-1}.s^{-1}$) | $5 \times 10^{-9} - 5 \times 10^{-6}$ | $0.16 \times 10^{-6}$ | $[-0.97, 30.63]$ | Parenchyma cells of corn leaves (20), *Chara* algae, onion and pea (21) |
| $\mu$ | Plasmodesmal permeability for open plasmodesmata ($m^3.Pa^{-1}.s^{-1}$) | $13.5 \times 10^{-27} - 0.33 \times 10^{-18}$ | $66.7 \times 10^{-24}$ | $[-1.00, 4947]$ | See Supplementary note |
| $\alpha$ | Source of solutes ($Pa.s^{-1}$) | $0.083 \times 10^3 - 0.167 \times 10^3$ | $0.118 \times 10^3$ | $[-0.30, 0.42]$ | Supplementary note and (11) |
| $\pi_{seed}$ | Osmotic pressure of seed proper ($Pa$) | $0.99 \times 10^6 - 1.29 \times 10^6$ | $1.13 \times 10^6$ | $[-0.12, 0.14]$ | Cotton (11), also see (22) |
| $P_{seed}$ | Turgor pressure of seed ($Pa$) | $0.07 \times 10^6 - 0.18 \times 10^6$ | $0.11 \times 10^6$ | $[-0.36, 0.64]$ | Cotton (11), also see (22) |
| $Y$ | Yield turgor pressure ($Pa$) | $0.06 \times 10^6 - 0.20 \times 10^6$ | $0.11 \times 10^6$ | $[-0.45, 0.82]$ | Assuming $P > Y$ and taking $P_{fiber}$ from (11) |
| $\phi$ | Cell Wall extensibility ($Pa^{-1}.s^{-1}$) | $1.1 \times 10^{-12} - 25.0 \times 10^{-12}$ | $5.3 \times 10^{-12}$ | $[-0.79, 3.72]$ | Growing pea stems (18, 23) |

**Fig. 2.**
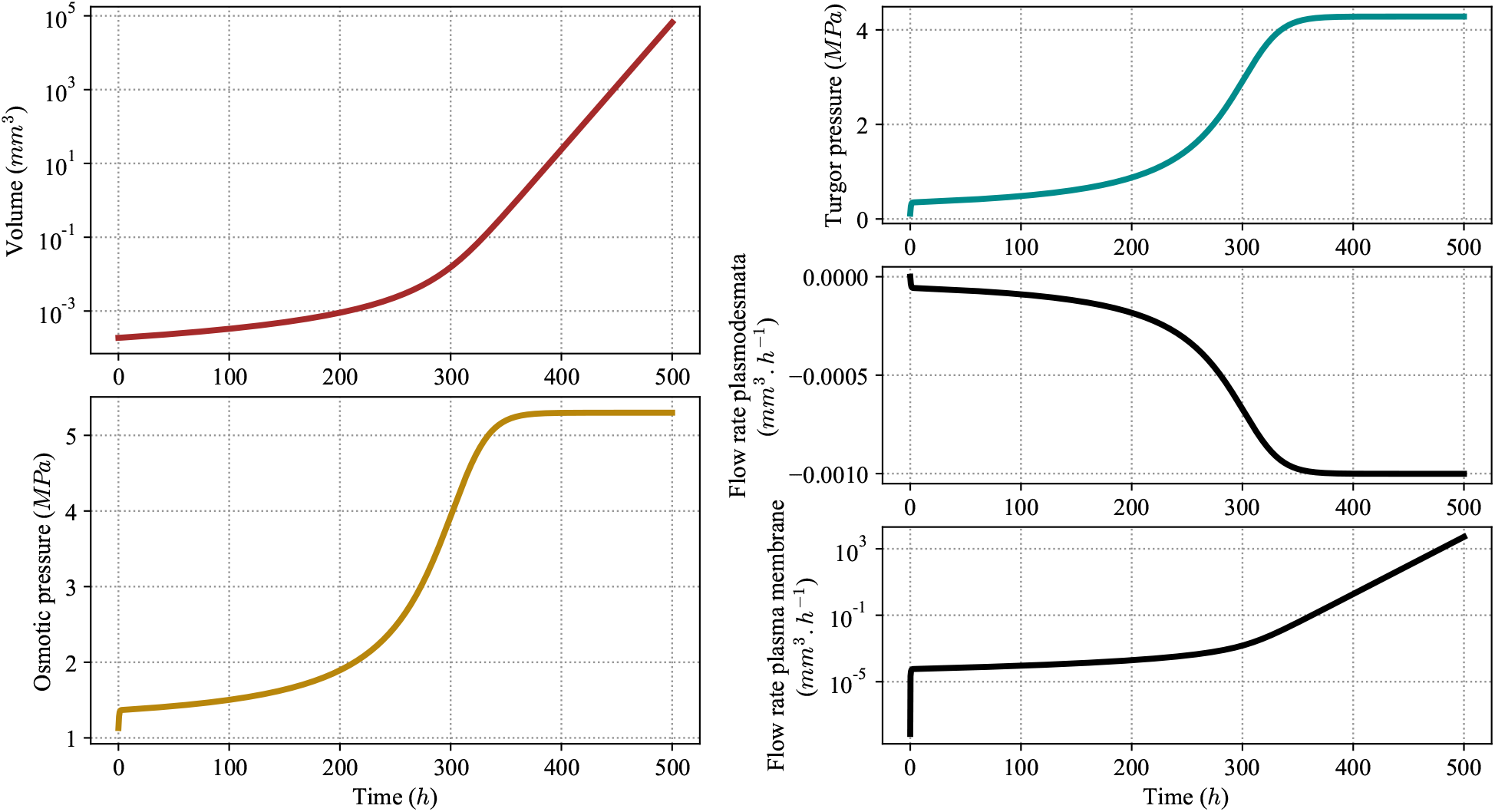
Behavior of the model with reference values of the parameters. Volume *V*, osmotic *π*_fiber_ and turgor *P*_fiber_ pressure, and flow rates through plasmodesmata *F*_*p*_ and through the plasma membrane *F*_*m*_, shown as a function of time. Reference values are given in Table 1. The initial value of the volume is *V* (0) = 1.88 *×* 10^*−*4^ *mm*^3^ and the initial value of the osmotic pressure is *π*_fiber_(0) = 1.13 *MPa*.

### Sensitivity of model predictions to parameter values

To identify the influence of parameters on the behavior of cotton fiber growth, we performed a one-factor-at-a-time sensitivity analysis of the reference model. To do this, we varied parameters one by one around their reference value by a maximal 10 percent deviation while keeping all others at their reference value, and we monitored the relative deviation of the final values of the three main observable variables: volume, osmotic pressure, and turgor pressure. As initial conditions for the differential equations, we assume that the turgor pressure in the fiber is initially higher than in the seed cells, as measured by (11, 22), while the duration of the simulation was chosen to be *t*_max_ = 500*h*, comparable with the natural duration of cotton fiber growth.

Detailed plots are shown in Figures S4-S6, while sensitivity values are given in Table 2. The sensitivity of an observable *X* to parameter *x* is defined as the normalized derivative of *X* with respect to *x* at its reference value,

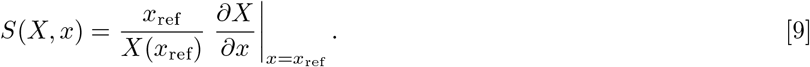

Table 2 presents the sensitivity of the logarithmic increase of volume log (*V* (*t*_max_)*/V* (0)), of the final osmotic pressure *π*_fiber_(*t*_max_), and of the final turgor pressure *P*_fiber_(*t*_max_) with to all the parameters of the model. As sensitivity values are normalized, they can be compared; their sign indicates positive or negative effect of the parameter on the observable. We see that some parameters potentially have a stronger influence on the final values of the volume and pressure than others (but some parameters have larger ranges of variation, see below).

**Table 2.** One-factor-at-a-time sensitivity around reference values. Values in the table are sensitivities *S*(*X, x*) of the observable *X* (rows) with respect to parameter *x* (columns). The final time *t*_max_ = 500*h* is comparable to the natural duration of cotton fiber growth.

| Variable | $L_r$ | $\mu$ | $\alpha$ | $\pi_{\text{seed}}$ | $P_{\text{seed}}$ | $Y$ | $\phi$ |
| --- | --- | --- | --- | --- | --- | --- | --- |
| Volume<br>$\log(V(t_{\max})/V(0))$ | $2.00 \times 10^{-4}$ | $-8.37 \times 10^{-1}$ | 1.35 | $-6.80 \times 10^{-1}$ | $2.79 \times 10^{-1}$ | $-2.74 \times 10^{-1}$ | 1.51 |
| Osmotic pressure<br>$\pi_{\text{fiber}}(t_{\max})$ | $-1.16 \times 10^{-5}$ | $-5.30 \times 10^{-6}$ | $4.41 \times 10^{-1}$ | $1.19 \times 10^{-1}$ | $-1.18 \times 10^{-2}$ | $1.16 \times 10^{-2}$ | $-4.42 \times 10^{-1}$ |
| Turgor pressure<br>$P_{\text{fiber}}(t_{\max})$ | $1.48 \times 10^{-5}$ | $-6.16 \times 10^{-6}$ | $5.46 \times 10^{-1}$ | $-1.16 \times 10^{-1}$ | $1.15 \times 10^{-2}$ | $1.44 \times 10^{-2}$ | $-5.47 \times 10^{-1}$ |

Regarding the effect on the final osmotic and turgor pressure values, we again observe two groups: *L*_*r*_, *P*_seed_ and *π*_seed_ have opposite effects on the two types of pressure, while theμ, *α, Y* and *φ* affect the two pressures in the same direction. Regarding the effect on growth, parameters can be classified into two groups: those that increase the final volume of the fiber – *L*_*r*_, *α, P*_seed_, and *φ* – and those that decrease it –μ, *π*_seed_, and *Y*. We also see that membrane permeability *L*_*r*_ has a lower effect on final size than the other parameters, while the closure of plasmodesmata as well as the increase of the solute source value have a higher growth-promoting effect.

### Effect of parameters on fiber length and comparison to experimental data

In what follows we further discuss how fiber length depends on parameters (see table 2 of sensitivities), and we show that the model is consistent with available experimental observations. We start with the normalized hydraulic conductivity of the cell membrane, *L*_*r*_. An increase in conductivity results in larger cell volume (the sensitivity is positive, though small) because it enables a higher influx of water for the same difference in water potential. Accordingly, two mutants with short fibres had lower aquaporin expression than in wild-type cotton (29). Moreover, downregulation of the aquaporin GhPIP2;6 led to shorter fibres (30). We nevertheless note that the predicted sensitivity of volume to hydraulic conductivity is low (of the order of 10^*−*4^) and that hydraulic conductivity was not quantified in the cotton fiber — for instance, the results of (29) might be ascribed to lower osmotic concentration in the fiber.

Plasmodesmatal permeablity,μ, has a negative effect on cell volume, because the fiber cell has higher turgor pressure than the seed coat and so may lose its contents through plasmodesmata. Consistently, the duration of plasmodesmata closure is correlated with fiber length across Gossypium species (12).

The solute source, *α*, has a positive effect on fiber volume. Indeed, accumulation of solutes increases fiber osmotic pressure, the driving force of growth. This is consistent with experiments that alter solute content of the fiber. Two mutants with short fibres had fiber cells with lower osmotic concentration (29) (though aquaporin expression was also altered in these mutants). When sucrose synthase is downregulated, sucrose synthase activity correlates well with fiber length (31). Finally, the vacuolar invertase GhVIN1 is highly expressed in fibers and its downregulation or upregulation respectively leads to shorter or to longer fibers (14).

Osmotic pressure of the seed coat, *π*_seed_, has a negative effect on fiber volume. Indeed, an increase in *π*_seed_ decreases the seed coat water potential and so reduces the relative advantage of the fiber regarding water potential, diminishing water flux in the fiber. Conversely, turgor pressure of the seed, *P*_seed_, has a positive effect on fiber volume because it contributes positively to water potential of the seed coat, in addition to negatively contributing to flow through plasmodesmata out of the fiber. These two parameters have not been manipulated experimentally (without affecting the fiber at the same time).

Yield threshold and extensibility have respectively negative and positive effect on fiber volume, as directly implied by Lockhart’s law (Eq. 7). Interestingly, GhTCP4 promotes secondary cell wall formation in cotton fibers(32), presumably reducing extensibility or increasing yield threshold. The downregulation or upregulation of GhTCP4 respectively leads to longer or to shorter fibers, consistent with such effect on extensibility or yield threshold.

### Model sensitivity over the biologically-relevant parameter range

The sensitivity analysis conducted above near the reference values was useful in unravelling how observable variables (notably fiber length) depend on parameters. However, that analysis implicitly assumed that model parameters have the same relative range of variations, which does not hold (as can be seen in Table 1). A parameter with broad biologically-relevant values may have a stronger impact on an observable variable than a parameter with narrow biological range, assuming similar sensitivity values of the observable variable to these two parameters. To overcome this shortcoming, we introduced the ‘maximal change of an observable variable *X* with respect to the parameter *x*’ and denote it *MC*(*X, x*). This quantity characterizes the effect of a parameter on the observable variables when it is allowed to sweep over the whole range of biologically-relevant values, we define it as

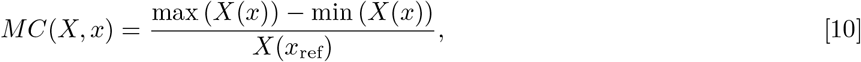

where the maximum and minimum of *X* are considered while *x ∈* [*x*_min_, *x*_max_] with *x*_min_ and *x*_max_ the biological limits of the parameter *x* in table 1, while keeping all other parameters at their reference value *x*_ref_. By definition, the sign of this maximal change is always positive.

We consider the same observable variables, as for the sensitivities in table 2, namely the logarithm of the ratio of final to initial volume, log (*V* (*t*_max_)*/V* (0)), the final osmotic pressure, *π*_fiber_(*t*_max_), and the final turgor pressure, *P*_fiber_(*t*_max_). We plotted the corresponding maximal changes on figure 3. These results single three parameters,μ, *α*, and *φ*, as the most influential on the observed final volume and pressures.

**Fig. 3.**
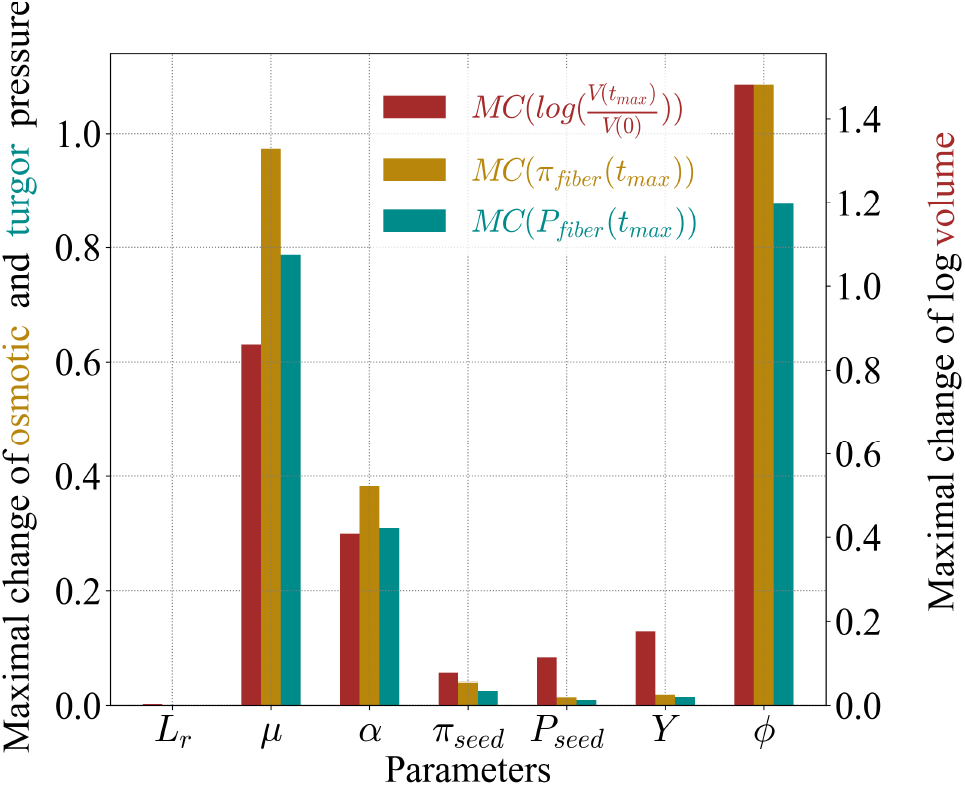
Maximal change. Plotted values are maximal changes *MC*(*X, x*) of the observable variable *X* (final volume, osmotic pressure, or turgor pressure) with respect to parameter *x* (hydraulic conductivity, plasmodesmatal permeablity, solute source, osmotic and turgor pressures of seed coat, yield threshold, and extensibility). This maximal change, defined in equation 10, takes into account the biological ranges of parameeters shown in Table 1. The final time *t*_max_ = 500*h* is comparable to the natural duration of cotton fiber growth.

### Constant parameters imply monotonic osmotic and turgor pressure

We now broadly analyse the behavior of the model, considering that the parameters take any value, and notably seek whether the model can predict the observed peaks in turgor and osmotic pressures (11). We sought a mathematical proof of possible model behaviors. To do so, we reduced the number of parameters from 7 physical parameters to 3 non-dimensional parameters by nondimensionalizing Equations (1), (6), and (8). We enumerated all possible behaviors and determined the range of parameter values that correspond to each type of behavior. The detailed proof, the list of all behavior types, and the associated conditions are presented in the Supplementary note.

Briefly, we proved that, depending on parameter values, we can have both monotonic and non-monotonic behaviors for turgor pressure and osmotic pressure. Having a peak of turgor or osmotic pressure is possible only when the source of solute *α* is strictly lower than a specific combination of the other parameters,

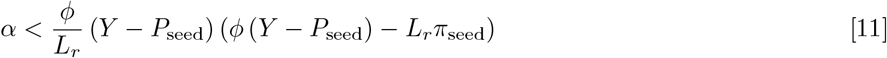

and when the turgor pressure in the fiber is strictly lower than in the seed. None of these two conditions is satisfied by experimental data. The biological range of Table 1 is incompatible with the first condition, while turgor pressure in fiber higher is higher than in the seed (11, 22).

Altogether, the model does not recover the observed non-monotonic behaviors of turgor and osmotic pressures (11) for constant parameters having values within the biologically-relevant ranges.

## A model with dynamic parameters retrieves experimental observations

Based on the above, we consider whether dynamic model parameters allow us to retrieve experimental observations; furthermore, it appears that three parameters *μ, α*, and *φ*, are those that have the highest impact on growth, and, if made variable in time, could potentially lead to a peak of osmotic and turgor pressures and to an arrest of growth.

### Vanishing cell wall extensibility yields growth arrest

We first address growth arrest, following Lockhart (1). Several pieces of evidence suggest a decrease in cell wall extensibility during fiber development. The expression of cell wall-related genes is dynamic (33), with notably a decrease of expression of genes involved in cell wall remodelling (33, 34) or an increase in expression of cellulose synthases associated with secondary cell wall (32). The relative quantity of cellulose, the stiffest component of the cell wall, increases during fiber development (34, 35), consistent with an increase in mechanical strength (36). Also, during fiber development, cellulose fibrils orientation shifts towards longitudinal (37), which effectively amounts to a decrease in extensibility in a one-dimensional model like ours (38). Accordingly, we considered that extensibility vanishes at 500 h and is a linear function of time taking its reference value at 0 h, as plotted in Figure 4-b)). The results are shown in figure 4-a). The volume reaches a constant value when the cell wall extensibility becomes zero. The final values of osmotic and turgor pressure are reduced with respect to the reference case.

**Fig. 4.**
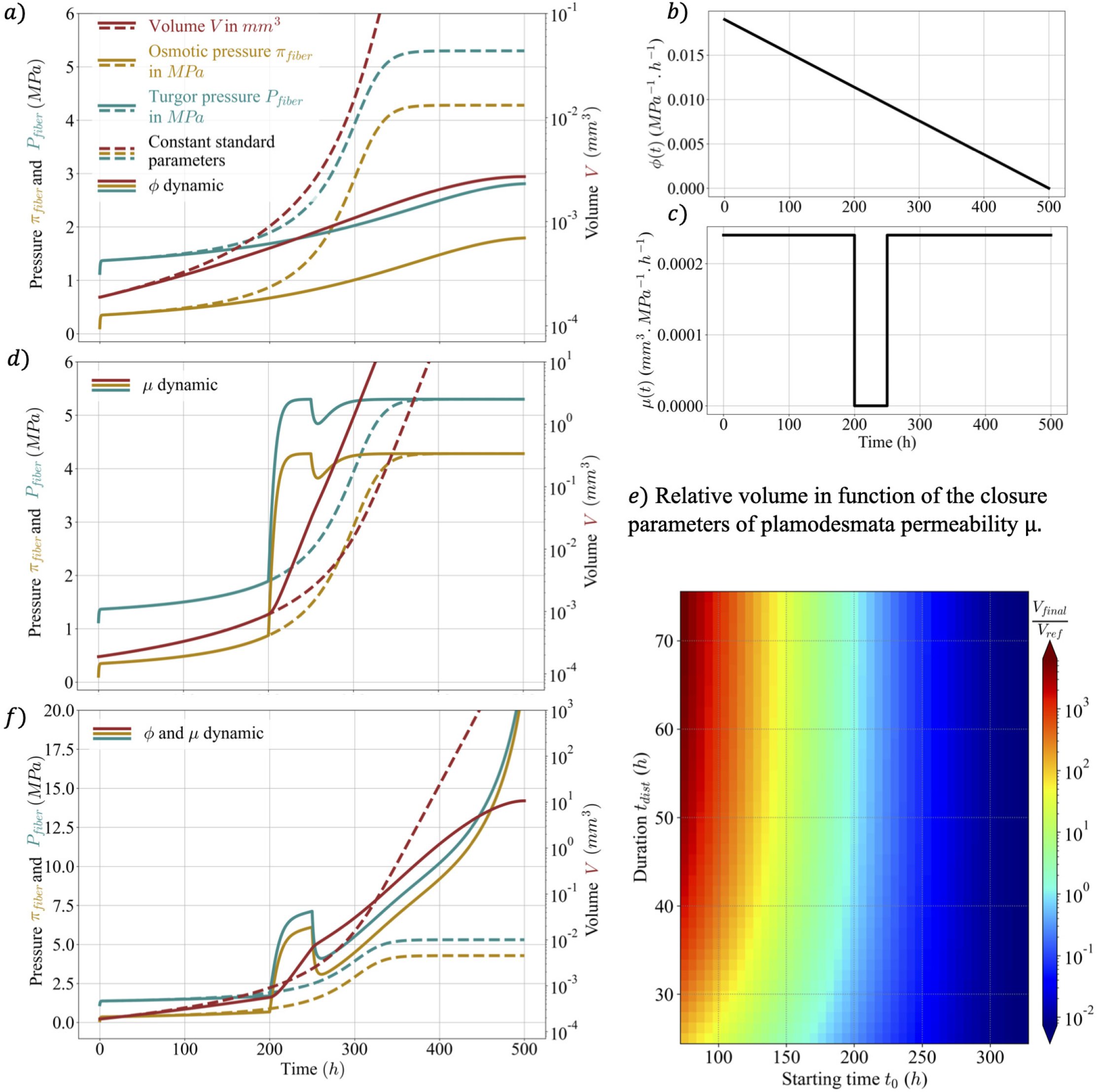
Predicted osmotic pressure, *π*_fiber_, turgor pressure, *P*_fiber_, and volume, *V*, of the fiber with dynamical parameters. In panels (a,d,f) the dashed lines correspond to model predictions when the parameters are constant at their reference values (same as in Fig. 2). In (a), continuous lines correspond to model predictions with dynamic extensibility, *φ*, following the temporal pattern shown in (b). In (d), continuous lines correspond to model predictions with dynamic plasmodesmata permeablity,μ, following the temporal pattern shown in (c). In (f), continuous lines correspond to model predictions with both *φ* andμ dynamic, combining the cases shown in (a) and (d). (e) Heat map of the normalised final volume of the fiber (colorscale on right) as as a function of the starting time, *t*_0_, and of the duration, *t*_*dist*_, of plasmodesmata closure. The final volume at 3000 h is normalized by its value for *t*_*dist*_ = 50 h and *t*_0_ = 200 h.

### Transient plasmodesmata closure or increased solute source yield a peak in the pressures

We now test the proposal by Ruan *et al*. (11) that plasmodesmata gating is needed for the peak in osmotic and turgor pressure. To do so, we consider that plasmodesmata permeability,μ, vanishes between 200 *h* and 250 *h*, as shown in Figure 4-b. This indeed induces a transient peak in turgor and osmotic pressure, see Figure 4-d). However, this peak is followed by a transient drop in pressure, which does not resemble experimental observations (11). As a consequence, additional hypotheses seem to be needed to fully explain the pressure behaviors in cotton fibers. When combining transient plasmodesmata gating and vanishing extensibility (Figure 4-f), the peak in pressures was more similar to experimental observations, although there was a later increase in pressure values.

We examined the effect on fiber volume of the temporal pattern of plasmodemata closure. In Figure 4-e), we plotted the ratio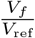as a function of the duration and the starting time of closure. This shows that longer fibers are favored by both earlier and longer plasmodemata gating. The former condition is consistent with experimental observations. Cotton lines with altered sterol levels had delayed plasmodesmata closure and shorter fibers (13).

We next consider the proposal by Ruan *et al*. (11) that an increase in solute content of the fiber is also needed for the peak in osmotic and turgor pressure. Several experimental results support such an increase in solute contents. There is peak in expression of genes related to osmolyte accumulation (39). Consistently, there is peak in the concentration of important osmolytes such as potassium/malate around 15DAA (15, 40). Accordingly, we considered a dynamic source of solute, with a transient rise from the reference value to the maximal biologically-relevant value (1) of the source parameter *α*. We observed a peak of turgor and osmotic pressure (Figure S7), but the height of the pressure peak was small, suggesting that solute dynamics has a minor role compared to plasmodesmata gating.

## Conclusions and perspectives

We have put forward a minimal differential equation-based model to explore the key ingredients for the striking elongation of cotton fibers. We notably assessed the contribution of plasmodesmal permeability in the behavior of turgor and osmotic pressures during cotton fiber development. To do so, we used Lockhart’s model in which we incorporated fluxes of water and solutes through plasmodesmata. Model predictions agreed with available experimental observations. We notably observed that a transient closure of plasmodesma increases fiber length and induces a peak in osmotic and turgor pressure, consistent with a qualitative model proposed based on experimental data (11).

Nevertheless, how this transient closure of plasmodesma is achieved is beyond the scope of this manuscript. It has been proposed that plasmodesmata may close when the difference in pressure between the two cells is high, through a valve-like mechanism (41, 42). This mechanism could explain the closing of the plasmodesmata during fiber development. However, when plasmodesmata are closed, the differences in turgor pressure between the fiber and the seed are increased and this mechanism would lead to the maintenance of closure. Another mechanism would be needed to explain the re-opening of plasmodesmata. It seems plausible that callose is involved in the control of plasmodesmata gating because callose deposition and degradation was observed to correlate respectively with closing and opening of plasmodesmata (12).

Our model has a few limitations. Although the shape of the fiber can be modelled (26), we only considered fiber length in our model, because it can be easily compared to experimental data and because it made it possible to broadly investigate the effect of parameter values on model predictions. We predicted plasma membrane conductivity to have a low (though positive) effect on fiber length, in contrast with experimental studies that associated lower aquaporin expression with significant reduction in fiber length (29, 30). Such discrepancy has also been noted in other contexts, questioning the validity of Lockhart’s model (43). The cotton fiber features a complex dynamics of solute import and conversion (e.g. by invertases) (14, 15, 31, 39, 40, 44). We considered solutes as whole and modelled a global source of solutes (parameter *α*) because we lack information to build a detailed model of solutes dynamics. Using such a parsimonious approach allowed us to qualitatively compare model predictions with experimental data. Regarding the transport of solutes, we only considered advection of solutes through plasmodesmata because when they are open, the difference in osmotic pressure between the fiber and the seed coat is very small (11), and so the differences in concentration of solutes are expected to be negligible, limiting the contribution of diffusion. Moreover, we considered a constant water potential all along the fiber, assuming that plasma membrane conductivity is limiting for water movement. Indeed, fibers are surrounded by water and they seem to lack cuticle during most of their development (45). Finally, we considered a single fiber, not accounting for the adhesion with neighboring fibers that occur during part of seed development (33, 45, 46). This is not a real limitation because adhesion would not affect mechanical stress patterns that drive growth.

Altogether, we found that a reduction in plasmodesmata permeability enables the building up of turgor pressure in a specific cell type, the cotton fiber. This phenomenon might have a broader relevance. The guard cells forming a stoma are symplasmically isolated from (are not connected with plasmodesmata to) neighboring cells, see e.g. (47), while they develop higher turgor pressure than their neighbors, see e.g. (48), which is essential to function of the stoma. Differences in turgor between cells within a tissue might also be important for cell size homeostasis (7, 49).

## Materials and Methods

Details are given in the supplementary note. Briefly, model parameters were determined based on a review of the literature, taken from cotton whenever available and from other species otherwise. Numerical solutions and graphs were produced with Python 3.9.5, notably using ‘odeint’ to solve differential equations. Formal analysis of model properties allowed us to conclude that the model predicts no pressure maxima.

## ACKNOWLEDGMENTS

This work was supported by CONACYT (scholarship #410744 to VHH) and Agence Nationale de la Recherche (project HydroField #ANR-20-CE13-0022-03 to AB). We are grateful to Mariana Benitez, to Yann Boursiac, to Ibrahim Cheddadi, and to Gywneth Ingram for their input into this project.

## Data Availability Statement

The authors confirm that the data used for this study are available within the article and its supplementary materials. References to the source publications for all measured or estimated parameter values are given (see Table 1).

## Notes

### Competing Interest Statement

The authors have declared no competing interest.

### Summary of Updates

Slight changes in the values of the reference parameters. These changes do not affect the conclusions in the former version. Clarifications were added in the main text. A study of the effect of the geometry on the permeability of plasmodesmata was added in the Supplementary materials.

