## Supplementary Material for "A mechanohydraulic model supports a role for plasmodesmata in cotton fiber elongation"

##### This PDF file includes:

Supplementary text
Figs. S1 to S7
Tables S1 to S4
SI References

### Supporting Information Text

**3 Summary of the analytical study and numerical illustrations** **15**

**4 Condition for a peak of turgor pressure and biologically-relevant parameters** **19**

**5 Details of the sensitivity analysis** **19**

**6 Dynamic source of solute** **21**

#### 1. Model parameters

Values for the model parameters listed in the main text in Table 1 are determined based on a literature review. In the absence in the literature of direct values for the plasmodesmal permeability and the solute source in cotton fibers, we estimated here these two parameter values, using other measurements from the literature. Parameter values used for these estimations together with the source references are summarised in Table S1. More precisely, for each parameter we identified a range of observed values, and we compute a corresponding reference value as the geometric mean of the range limits. We also verified that the typical values used in other examples of modeling approaches to plasmodesmal permeability, like in (1, 2), lie within the range that we used.

**A. Estimation of plasmodesmal permeability.** To estimate plasmodesmal permeability $\mu$, we followed the methodology of (3) and first calculated the conductivity of a single plasmodesma using different laws depending on the assumptions that we made, see Fig. S1.

If a plasmodesma were just an empty cylindrical tube of radius $R$ and length $w$, its permeability $\lambda_R$ to a fluid of viscosity $\eta$ would be given by the Hagen-Poiseuille law,

$$\lambda_R = \frac{\pi R^4}{8\eta w}. \quad [1]$$

However, the desmotubule obstructs the middle of the plasmodesma and can be considered as a cylinder of radius $\gamma R$ with $\gamma \in [0, 1]$. Therefore, even if we assume that the space between the desmotubule and the plasma membrane (cytoplasmic sleeve) is free, the Hagen-Poiseuille law has to be modified to account for the occlusion of the plasmodesma channels by the desmotubule, see

Figures S1-a for the transversal and S1-b for the longitudinal section. In this case, the equation for annular flow yields the permeability of a single plasmodesma, (1, 4, 5)

$$\lambda_{central} = \lambda_R \left(1-\gamma^2\right) \left[1+\gamma^2+\frac{1-\gamma^2}{\ln(\gamma)}\right]. \quad [2]$$

We plotted the dependence of this permeability on the relative desmotubule radius $\gamma$ in Figure S1-g. Notice that the permeability drastically drops with the presence of the desmotubule: it loses one order of magnitude with respect to the empty cylinder with a relative desmotubule radius of 0.5. For desmotubules that are not very thin, this permeability can be approximated quite well (see Figure S1-g) by the following polynomial expression in $\gamma$

$$\lambda_{central\_proxy} = \frac{2}{3}\lambda_R \left(1+\gamma\right)\left(1-\gamma\right)^3. \quad [3]$$

If we consider now that the space between the desmotubule and the plasma membrane is obstructed so that the cylindrical channels of radius $k$ in the ultrastructure of the plasmodesma take half of the annular volume left by the desmotubule (6–8), see Figure S1-c, then the permeability of a single plasmodesma becomes

$$\lambda_{obstructed} = \lambda_R \left(1-\gamma^2\right) \frac{k^2}{2R^2}. \quad [4]$$

Notice that the channels have to fit in the cytoplasmic sleeve, therefore, they have a maximal radius $k_{max} = R(1-\gamma)/2$, given the plasmodesma and desmotubule radii. In Figure S1-g, we represented the permeability $\lambda_{obstructed}$ as a function of $\gamma$ with the reference value for $k$ based on measurements (see Table S1), unless this exceeds the maximal value $k_{max}$ for a given $\gamma$ value, where we considered $k = k_{max}$ instead.

Figure S1-g allows us to compare the predictions for the permeability of a single cylindrical plasmodesma given the above-presented models: central desmotubule with and without obstructed cytoplasmic sleeve. In particular, for the same radius $R$, length $w$, and relative desmotubule radius $\gamma$, we can get an upper estimation for the permeability by considering the empty cytoplasmic sleeve, as well as a lower estimation by considering an obstructed plasmodesma with channels. Therefore, by considering the whole biologically relevant parameter space (see Table S1) for the parameters appearing in these two models (central and obstructed), and taking the minimal and the maximal plasmodesmata permeability value, we estimate a range for the plasmodesmata permeability $\lambda$.

We note that other studies have explored more complex geometries, such as a desmotubule with an offset or varying radius (2), necks (9) or funnel-shaped plasmodesmata (10). Such geometries lead to changes in the permeability of a plasmodesma with respect to the above-presented geometries based on an ideal cylinder. In the following, we briefly present some of these more complex geometries as perturbations of the cylindric one, so that we can estimate the effect of these geometrical perturbations on the permeability of one plasmodesma.

On the one hand, in (2), they study for instance the effect of a possible offset of the desmotubule inside the plasmodesma. Let $\delta \in [0,1]$ be the offset parameter, namely the distance between the axes of the desmotubule and of the plasmodesma is $\delta(1-\gamma)R$, as schematized in S1-c. Then the permeability of a single plasmodesma with a relatively wide desmotubule increases quadratically with the offset parameter $\delta$

$$\lambda_{offset}(\delta) = \lambda_{central}\left(1+\frac{3}{2}\delta^2\right). \quad [5]$$

In particular, the permeability increases as soon as there is an offset of the desmotubule, and it can increase by a maximal value of 150% with respect to the unperturbed concentric plasmodesma when the desmotubule is in contact with the plasma membrane ($\delta = 1$). We traced the permeability $\lambda_{offset}(\delta = 1)$ in this extreme situation in figure S1-h.

On the other hand, a neck reduces permeability in the same proportions as an obstruction. To see this, we consider a plasmodesma with a neck region as presented in figure S1-e. In this context, the neck represents an annular obstruction of thickness $\omega(1-\gamma)R$ with $\omega \in [0,1]$ and length $\zeta w$ with $\zeta \in [0,1]$. This structure can be seen as a series of two annular pipes with different outer radii, and therefore, we approximate its permeability by

$$\lambda_{neck}(\omega,\zeta) = \frac{2}{3}\lambda_R \frac{1}{\frac{\zeta}{(1+\gamma-\omega(1-\gamma))(1-\gamma-\omega(1-\gamma))^3} + \frac{1-\zeta}{(1+\gamma)(1-\gamma)^3}}. \quad [6]$$

We represented in figure S1-h the permeability corresponding to this model taking into account a neck region which obstructs 50% of the cytoplasmic sleeve thickness ($\omega = 0.5$), on 10% of the length of the plasmodesma ($\zeta = 0.1$). One can see that indeed, the presence of the neck region reduces the permeability of the plasmodesma, and its effect is of the same order of magnitude as of the obstruction in the obstructed model.

Furthermore, the widening on one end of funnel-shaped plasmodesmata can drastically raise their permeability. Considering a plasmodesma of radius $R$ on the one end and of radius $(1+\epsilon)R$ on the other end, with a concentric desmotubule of radius $\gamma R$, as shown in figure S1-f, we obtain the following expression for its permeability

$$\lambda_{funnel}(\epsilon) = \frac{16}{3}\lambda_R \frac{\epsilon\gamma^3}{2\gamma\left(\frac{1-2\gamma+\epsilon}{(1-\gamma+\epsilon)^2} - \frac{1-2\gamma}{(1-\gamma)^2}\right) + \ln\left|\frac{1-\gamma+\epsilon}{(1+\gamma+\epsilon)^2}\right| - \ln\left|\frac{1-\gamma}{(1+\gamma)^2}\right|}. \quad [7]$$

We represented in figure S1-h this function of the relative desmotubule radius $\gamma$ for the specific values of $\epsilon = 0.5$ for a 50% and $\epsilon = 1$ for a 100% raising of the plasmodesmata radius on the one end. We see that an order of magnitude in permeability can easily be gained from this type of widening. However, since we did not find in the literature evidence for the existence of these funnel-shaped plasmodesmata for cotton on the cell wall separating the seed and the fiber, we did not consider this model in the estimation of the biological range of plasmodesma permeability.

Finally, the value of the conductivity of a plasmodesma, $\lambda$, was then scaled to the cell by multiplying it by the surface area of the cell wall shared by the fiber and the contiguous epidermal cell, and $\rho$ the surface density of the plasmodesmata. This yields the plasmodesmal permeability of the fiber,

$$\mu = \lambda\rho\Pi r^2 \quad [8]$$

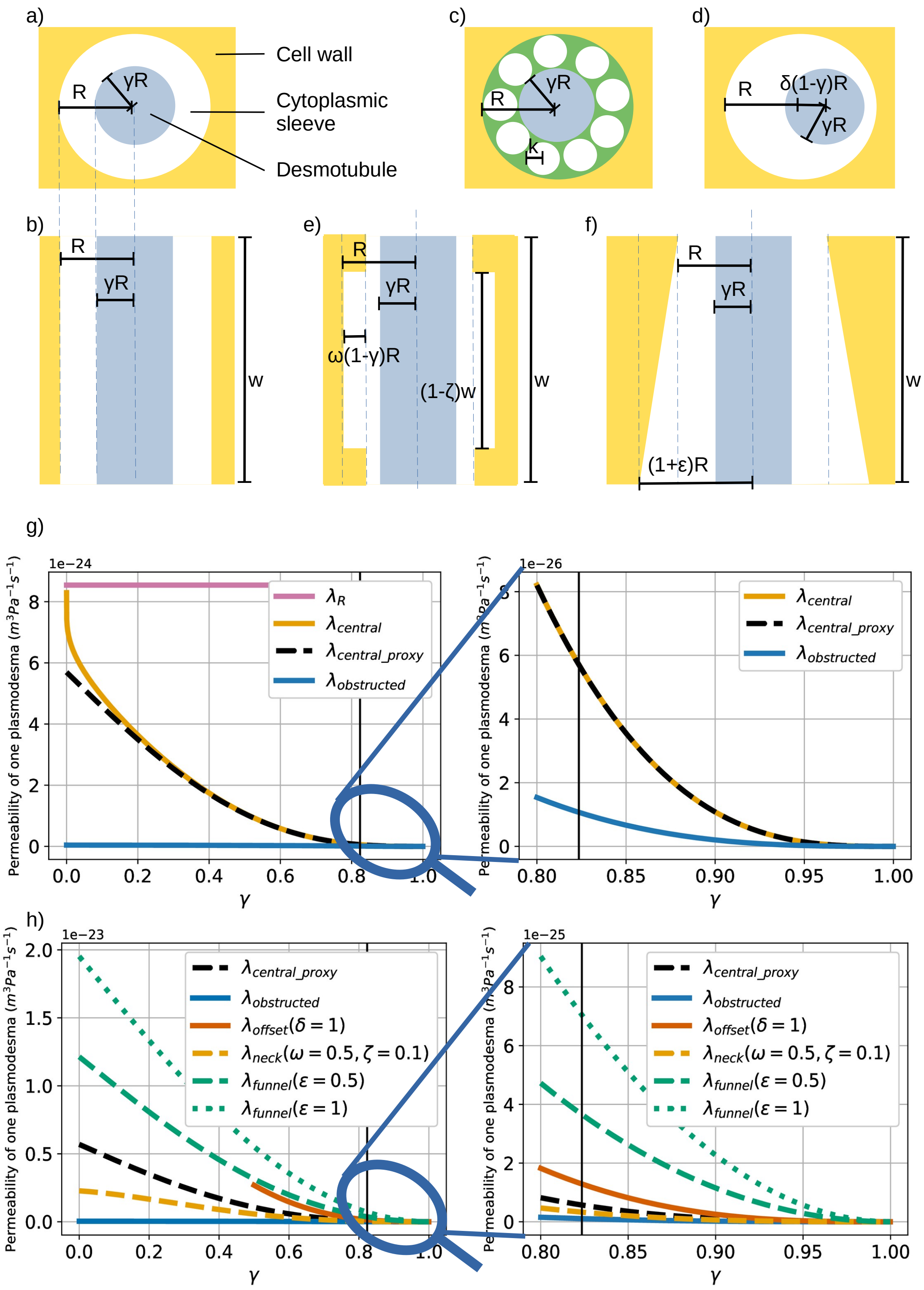

**Fig. S1.** **The effect of the geometry on the permeability of plasmodesmata.** Different models of a plasmodesma are presented as follows: a) transversal section of a plasmodesma with a concentric desmotubule; b) longitudinal section of a cylindrical plasmodesma with concentrical desmotubule; c) transversal section of an obstructed plasmodesma; d) transversal section of a plasmodesma with a desmotubule with an offset; e) longitudinal section when necks obstruct the plasmodesma; f) longitudinal section of funnel shaped plasmodesmata. g) Permeability $\lambda$ as a function of the relative desmotubule radius $\gamma$. The effect of an offset desmotubule and of the obstruction of the cytoplasmic sleeve in the case of a cylindrical plasmodesma are shown. h) The effect of necks and of a funnel shape are compared to the effect of the preceding perturbations. For g) and h) unless explicitly mentioned, all parameter values take the reference values from Table S1. The reference value for the relative desmotubule radius ($\gamma = 0.82$) is indicated by a vertical black line on the graphics.

**Table S1. List of the biological parameters for plasmodesmata.**

| Symbol | Name | Range | Reference value | Reference and notes |
|---|---|---|---|---|
| $w$ | Plasmodesma length $(m)$ | $0.23 \times 10^{-6} - 6 \times 10^{-6}$ | $1.2 \times 10^{-6}$ | Extrafloral nectaries of *G. hirsutum* (11), wall thickness variation from the start to the end of G. hirsutum fiber growth (12) |
| $R$ | Plasmodesma radius $(m)$ | $10 \times 10^{-9} - 30 \times 10^{-9}$ | $17 \times 10^{-9}$ | Extrafloral nectaries of *G. hirsutum* (11) |
| $\gamma R$ | Desmotubule radius $(m)$ | $9.2 \times 10^{-9} - 21 \times 10^{-9}$ | $14 \times 10^{-9}$ | Extrafloral nectaries of *G. hirsutum* (11) |
| $k$ | Radius of channel through plasmodesma ultrastructure $(m)$ | $1 \times 10^{-9} - 3 \times 10^{-9}$ | $1.7 \times 10^{-9}$ | Values from (3, 7) |
| $\rho$ | Plasmodesmata density $(m^{-2})$ | $0.5 \times 10^{12} - 12.4 \times 10^{12}$ | $2.5 \times 10^{12}$ | Minimal value: tobacco trichomes (13). Maximal value: extrafloral nectaries of *G. hirsutum* (11); |
| $r$ | Fiber radius $(m)$ | $10 \times 10^{-6} - 20 \times 10^{-6}$ | $14 \times 10^{-6}$ | *G hirsutum* fiber cells (14, 15) |
| $\eta$ | Cell sap dynamic viscosity $(Pa.s)$ | $2 \times 10^{-3} - 5 \times 10^{-3}$ | $3.2 \times 10^{-3}$ | Phloem cells (5, 16) |
| $\mu$ | Total plasmodesmal permeability for open plasmodesmata $(m^3 Pa^{-1} s^{-1})$ | $13.5 \times 10^{-27} - 0.33 \times 10^{-18}$ | $66.7 \times 10^{-24}$ | Calculated with equation (8), geometric mean of the maximum and minimum permeability values enabled by the parameter's range. |

Recall, that in order to estimate the range for the total plasmodesmal permeability $\mu$ in Table S1, we considered all the parameter space defined by the ranges for the other parameters in this table, considered the models of non-obstructed (equation 2) and obstructed (equation 4) cylindrical plasmodesma with concentrical desmotubule, and took the minimal and maximal value for $\mu$. We note that if we would have taken just the reference value of the parameters to estimate the range of $\mu$, then we would have obtained the narrow interval $21.2 \times 10^{-24} - 87.9 \times 10^{-24} m^3 Pa^{-1} s^{-1}$, and a reference value of $43.1 \times 10^{-24} m^3 Pa^{-1} s^{-1}$ which is very close to the value obtained in Table S1, computed using the whole parameter space.

**B. Estimation of the source of solute.** Since we assumed that $\pi_{\text{fiber}}$ is proportional to the concentration of solutes, we estimated the typical value of the source parameter, $\alpha$, as $\pi_{\text{fiber}}\kappa$, where $\pi_{\text{fiber}}$ was extracted from (17) and $\kappa$ is the average value of growth rate that we estimated from the increase of fiber length with time. We extracted experimental data of the length of cotton fiber cells reported in (18–21). We fitted lengths to the logistic growth equation:

$$f(t) = \frac{M}{1 + e^{-\kappa(t - t_0)}}, \qquad [9]$$

where $M$ is the maximum volume and $t_0$ is the midpoint of the curve. We then used the average value of $\kappa$.

#### 2. Analytical study of the behaviors of the system

Here we aim to mathematically address whether the model with constant parameters can predict a peak in (turgor or osmotic) pressure. To do so, we non-dimensionalized the governing equations and we studied the sign of the second derivative of the turgor and osmotic pressure with respect to time. We considered all possible cases, according to dimensionless parameters and we analytically derived conditions for the existence of a peak in pressure. This section contains the derivation of the equations to identify the different possible behaviors of the system. The main question is to know whether the model can have non-monotonic behaviors for the turgor and osmotic pressure like reported in experimental data.

Throughout this analysis we use the following dimensionless form of the equations of the model

$$\frac{\mathrm{d}v}{\mathrm{d}\tau} = v\pi - (v+1)\,p, \tag{10}$$

$$\frac{\mathrm{d}\pi}{\mathrm{d}\tau} = \alpha_0 - \frac{\pi}{v}(-p)_+ - (\pi - p)(\pi + 1), \tag{11}$$

$$p = \begin{cases} \dfrac{v\pi}{v+1} \text{ if } p \le y_0, \\ \\ \dfrac{v(\pi + \phi_0 y_0)}{v(1+\phi_0)+1} \text{ else.} \end{cases} \tag{12}$$

The dimensionless variables are defined from turgor pressure, osmotic pressure, cell volume, as

$$p = \frac{P_{\text{fiber}} - P_{\text{seed}}}{\pi_{\text{seed}}}, \tag{13}$$

$$\pi = \frac{\pi_{\text{fiber}} - \pi_{\text{seed}}}{\pi_{\text{seed}}}, \tag{14}$$

$$v = \frac{VL_r}{\mu}, \tag{15}$$

$$\tau = L_r \pi_{\text{seed}}^2 t. \tag{16}$$

And the dimensionless parameters are defined by

$$\alpha_0 = \frac{\alpha}{L_r \pi_{\text{seed}}^2}, \tag{17}$$

$$\phi_0 = \frac{\phi}{L_r}, \tag{18}$$

$$y_0 = \frac{Y - P_{\text{seed}}}{\pi_{\text{seed}}}. \tag{19}$$

**Table S2. List of the range of the dimensionless parameters with parameters values of table 1.**

| Symbol | Expression | Range |
|---|---|---|
| $\alpha_0$ | $\frac{\alpha}{L_r \pi_{\text{seed}}^2}$ | $[1.0 \times 10^{-5}, 3.4 \times 10^{-2}]$ |
| $\phi_0$ | $\frac{\phi}{L_r}$ | $[2.2 \times 10^{-7}, 5.0 \times 10^{-3}]$ |
| $y_0$ | $\frac{Y - P_{\text{seed}}}{\pi_{\text{seed}}}$ | $[-0.12, 0.13]$ |

To make it easier for the reader, we only provide here the analysis of the behavior of $p$. We note that we obtained the same results for $\pi$.

**A. Case without growth.** When turgor pressure in the fiber $P_{\text{fiber}}$ is lower than the yield threshold $Y$, we get

$$p = \frac{v\pi}{v+1}.$$

Replacing in equation (10) we obtain for the volume:

$$\frac{\mathrm{d}v}{\mathrm{d}\tau} = 0.$$

The volume remains constant, which closes this case.

**B. Case with growth.** Throughout this part, we assume that turgor pressure in the fiber $P_{\text{fiber}}$ is higher than the yield threshold $Y$ (in terms of the dimensionless parameters $p > y_0$). We consider the equations (10) and (11) and the dimensionless turgor pressure $p$ is

$$p = \frac{v\left(\pi + \phi_0 y_0\right)}{v\left(1+\phi_0\right)+1}. \qquad [20]$$

It is easy to check that the first derivative of $v$ is always positive if $p > y_0$. From Eqs. (20) and (10), we obtain

$$\frac{\mathrm{d}v}{\mathrm{d}\tau} = v\phi_0\left(p - y_0\right) > 0. \qquad [21]$$

The first derivative of the dimensionless turgor pressure, $p$, is

$$\frac{\mathrm{d}p}{\mathrm{d}\tau} = \frac{v}{v\left(1+\phi_0\right)+1}\left(\frac{\mathrm{d}\pi}{\mathrm{d}\tau} + \frac{p}{v^2}\frac{\mathrm{d}v}{\mathrm{d}\tau}\right). \qquad [22]$$

Because Eq. (11) contains the positive part of p we distinguish the cases when $p$ is strictly negative (section B.1) and when $p$ is positive or null (section B.2).

First of all, we write below the second derivative of the dimensionless volume, $v$, the dimensionless osmotic pressure, $\pi$, and the dimensionless turgor pressure, $p$, that we will use in the following sections. Equation (10)) yields

$$\frac{\mathrm{d}^2v}{\mathrm{d}\tau^2} = v\frac{\mathrm{d}\pi}{\mathrm{d}\tau} + \left(\pi - p\right)\frac{\mathrm{d}v}{\mathrm{d}\tau} - \left(v+1\right)\frac{\mathrm{d}p}{\mathrm{d}\tau}. \qquad [23]$$

Equation (11) yields

$$\frac{\mathrm{d}^2\pi}{\mathrm{d}\tau^2} = -\frac{1}{v}\left(-p\right)_+\frac{\mathrm{d}\pi}{\mathrm{d}\tau} + \frac{\pi}{v^2}\left(-p\right)_+\frac{\mathrm{d}v}{\mathrm{d}\tau} - \frac{\pi}{v}\frac{\mathrm{d}}{\mathrm{d}\tau}\left(-p\right)_+ - \left(\pi - p\right)\frac{\mathrm{d}\pi}{\mathrm{d}\tau} - \left(\pi+1\right)\left(\frac{\mathrm{d}\pi}{\mathrm{d}\tau} - \frac{\mathrm{d}p}{\mathrm{d}\tau}\right). \qquad [24]$$

Finally, Equation (22) results in

$$\frac{\mathrm{d}^2p}{\mathrm{d}\tau^2} = \frac{1}{1+v\left(1+\phi_0\right)}\left(v\frac{\mathrm{d}^2\pi}{\mathrm{d}\tau^2} + \frac{p}{v}\frac{\mathrm{d}^2v}{\mathrm{d}\tau^2} + \frac{2}{v}\frac{\mathrm{d}p}{\mathrm{d}\tau}\frac{\mathrm{d}v}{\mathrm{d}\tau} - \frac{2p}{v^2}\left(\frac{\mathrm{d}v}{\mathrm{d}\tau}\right)^2\right). \qquad [25]$$

In this analysis, we seek whether $p$ reaches a maximum. We thus assume the existence of a dimensionless time $\tau_0 \geq 0$ such that the first derivative of the dimensionless turgor pressure $p$ is equal to zero

$$\frac{\mathrm{d}p}{\mathrm{d}\tau}\left(\tau_0\right) = 0.$$

This can be rewritten using Eq. (22) as

$$\frac{\mathrm{d}\pi}{\mathrm{d}\tau}(\tau_0) + \frac{p}{v^2}\frac{\mathrm{d}v}{\mathrm{d}\tau}(\tau_0) = 0. \qquad [26]$$

Then, using the Eq. (25), we find a necessary condition to have the second derivative of the dimensionless turgor pressure $p$ negative

$$\frac{\mathrm{d}^2 p}{\mathrm{d}\tau^2}(\tau_0) < 0 \implies v\frac{\mathrm{d}^2\pi}{\mathrm{d}\tau^2}(\tau_0) + \frac{p}{v}\frac{\mathrm{d}^2 v}{\mathrm{d}\tau^2}(\tau_0) - \frac{2p}{v^2}\left(\frac{\mathrm{d}v}{\mathrm{d}\tau}(\tau_0)\right)^2 < 0.$$

Hereafter we consider $p$ and other quantities to be evaluated at time $\tau_0$, dropping $(\tau_0)$ for brievity. By developing the equation (26) using the differential equations (10) and (11) and the expression of $p$ (20) we obtain

$$\begin{aligned}\frac{\mathrm{d}p}{\mathrm{d}\tau}(\tau_0) = 0 &\implies \frac{\mathrm{d}\pi}{\mathrm{d}\tau}(\tau_0) + \frac{p}{v^2}\frac{\mathrm{d}v}{\mathrm{d}\tau}(\tau_0) = 0,\\ &\implies -p^2\left[1 + v\left(1 + v\phi_0\right)\left(1 + \phi_0\right)\right] + pv\left[\phi_0 y_0 v\left(1 + \phi_0\right) + \left(1 + v\phi_0\right)\left(\phi_0 y_0 - 1\right)\right]\\ &\quad + \left(v\phi_0 y_0 - p\left(v + \phi_0 v + 1\right)\right)\left(-p\right)_+ + v^2\left(\alpha_0 + \phi_0 y_0\left(1 - \phi_0 y_0\right)\right) = 0. \end{aligned} \qquad [27]$$

We obtain therefore a quadratic polynomial of $p$. This polynomial can have either zero, one or two solutions corresponding to the value of the dimensionless turgor pressure $p$ at dimensionless time $\tau_0$ when the first derivative of $p$ equals zero. We study then independently the different cases depending on the sign of $p$.

***B.1. Case $P_{fiber} < P_{seed}$ ($p < 0$).*** We seek to demonstrate in this section that the dimensionless turgor pressure $p \mapsto p(\tau)$ defined over $\mathbb{R}_+$ can have at least one local maximum, implying that the function $p$ can be non-monotonic with a local maximum. Because $p < 0$ in this section, the dimensionless yield threshold $y_0$ must be strictly negative, $y_0 < 0$. We reformulate Eq. (27) using $p < 0$. At $\tau = \tau_0$ we have

$$\begin{aligned}\frac{\mathrm{d}p}{\mathrm{d}\tau}(\tau_0) = 0 \implies &-p^2\left[v^2\phi_0\left(1 + \phi_0\right)\right] + pv\left[\phi_0 y_0\left(v + v\phi_0 - 1\right) + \left(1 + v\phi_0\right)\left(\phi_0 y_0 - 1\right)\right]\\ &+ v^2\left(\alpha_0 + \phi_0 y_0\left(1 - \phi_0 y_0\right)\right) = 0.\end{aligned} \qquad [28]$$

The discriminant of this quadratic polynomial is

$$\Delta = v^2\left[\left[\phi_0 y_0\left(v + v\phi_0 - 1\right) + \left(1 + v\phi_0\right)\left(\phi_0 y_0 - 1\right)\right]^2 + 4v^2\left[\alpha_0 + \phi_0 y_0\left(1 - \phi_0 y_0\right)\right]\left[\phi_0\left(1 + \phi_0\right)\right]\right].$$

Let us study the sign of the discriminant $\Delta$. The discriminant is positive if and only if

$$v^2\left[4\alpha_0\phi_0\left(1 + \phi_0\right) + \phi_0^2\left(1 + y_0\right)^2\right] + 2v\phi_0\left[1 - y_0\left(1 + 2\phi_0\right)\right] + 1 \geq 0. \qquad [29]$$

Since $y_0 < 0$, all terms of this quadratic polynomial of $v$ (in Eq. 29) are strictly positive. Therefore $\Delta > 0$ and Eq. (28) has two solutions $p_+$ and $p_-$ such that

$$p_\pm = \frac{1}{2v^2}\frac{v\phi_0 y_0\left(v + v\phi_0 - 1\right) + v\left(1 + v\phi_0\right)\left(\phi_0 y_0 - 1\right) \pm \sqrt{\Delta}}{\phi_0\left(1 + \phi_0\right)}.$$

As in this section $p(\tau_0) > y_0$ and $p(\tau_0) < 0$, we must verify whether the solutions $p_+$ and $p_-$ satisfy these conditions. Let us start with the condition $p(\tau_0) < 0$. We have

$$p_\pm < 0 \iff \pm\sqrt{\Delta} < q(v) \text{ with } q(v) = -v\phi_0 y_0 \left(v\left(1+\phi_0\right)-1\right) - v\left(1+v\phi_0\right)\left(\phi_0 y_0 - 1\right). \qquad [30]$$

In fact $q(v) > 0$ because,

$$q(v) > 0 \iff v\phi_0 \left[y_0\left(2\phi_0+1\right)-1\right] < 1.$$

and $y_0 < 0$, $v > 0$ and $\phi_0 > 0$. As a consequence, $p_-$ always satisfies Eq. (30). We therefore focus on the solution $p_+$. Eq. (30) is then equivalent to

$$\begin{aligned} &\Delta < v^2\left[\phi_0 y_0\left(v+v\phi_0-1\right)+\left(1+v\phi_0\right)\left(\phi_0 y_0-1\right)\right]^2, \\ \iff &\left[\alpha_0+\phi_0 y_0\left(1-\phi_0 y_0\right)\right]\left[\phi_0\left(1+\phi_0\right)\right] < 0, \\ \iff &\alpha_0 < \phi_0 y_0\left(\phi_0 y_0 - 1\right). \end{aligned}$$

If this condition $\alpha_0 < \phi_0 y_0 (\phi_0 y_0 - 1)$ is verified, then the solution $p_+$, as well as $p_-$, satisfies the conditions $\frac{\mathrm{d}p}{\mathrm{d}\tau}(\tau_0) = 0$ and $p(\tau_0) < 0$. Otherwise, if $\alpha_0 \geq \phi_0 y_0 (\phi_0 y_0 - 1)$, the solution $p_+$ does not satisfy the condition $p(\tau_0) < 0$, then we have a single solution, $p_-$.

Next we consider the condition $p(\tau_0) > y_0$. We have

$$p_-(\tau_0) > y_0 \iff \sqrt{\Delta} < -v\left(1+v\phi_0\left(1+y_0\right)\right), \qquad [31]$$

and

$$p_+(\tau_0) > y_0 \iff \sqrt{\Delta} > v\left(1+v\phi_0\left(1+y_0\right)\right). \qquad [32]$$

In Eq. (31), $p_-$ does not satisfy the condition $p_-(\tau_0) > y_0$ if the quantity $1 + v\phi_0(1+y_0)$ is positive. If this last quantity is negative, then after calculating we find

$$p_-(\tau_0) > y_0 \iff v\alpha_0 < y_0,$$

which is false, because $\alpha_0 > 0$, $v > 0$, and we assumed that $y_0 < 0$. Therefore $p_-$ does not meet the condition $p_-(\tau_0) > y_0$, and is not a possible solution for $\frac{\mathrm{d}p}{\mathrm{d}\tau}(\tau_0) = 0$ and $p(\tau_0) < 0$.

In Eq. (32), $p_+$ satisfies $p_+(\tau_0) > y_0$ if the quantity $v(1 + v\phi_0(1+y_0))$ (which implies $y_0 < -1$) is negative. If this last quantity is positive, then Eq. (32) can be shown to be equivalent to

$$\Delta > v^2\left(1+v\phi_0\left(1+y_0\right)\right)^2 \iff v\alpha_0 > y_0,$$

which is true because $\alpha_0 > 0$, $v > 0$, and we assumed that $y_0 < 0$. We conclude that we have $p_+(\tau_0) > y_0$ without additional conditions on the parameters.

So far, we have analyzed whether the first derivative of $p(\tau)$ vanishes. We found that the only solution satisfying $p(\tau_0) < 0$ and $p(\tau_0) > y_0$ is $p_+$. We next study the sign of the second derivative of the dimensionless turgor pressure $p$. As we assume that $p < 0$, Eq. (24) becomes

$$\frac{\mathrm{d}^2\pi}{\mathrm{d}\tau^2} = \frac{p}{v}\frac{\mathrm{d}\pi}{\mathrm{d}\tau} - \frac{\pi p}{v^2}\frac{\mathrm{d}v}{\mathrm{d}\tau} - \frac{\pi}{v}\frac{\mathrm{d}p}{\mathrm{d}\tau} - (\pi - p)\frac{\mathrm{d}\pi}{\mathrm{d}\tau} - (\pi+1)\left(\frac{\mathrm{d}\pi}{\mathrm{d}\tau} - \frac{\mathrm{d}p}{\mathrm{d}\tau}\right).$$

For $\tau = \tau_0$ (implying that $\mathrm{d}p/\mathrm{d}\tau = 0$), this yields

$$\frac{\mathrm{d}^2\pi}{\mathrm{d}\tau^2}(\tau_0) = \frac{\mathrm{d}\pi}{\mathrm{d}\tau}(\tau_0)\left(\frac{p}{v} + p - 2\pi - 1\right) - \frac{\pi p}{v^2}\frac{\mathrm{d}v}{\mathrm{d}\tau}(\tau_0).$$

Meanwhile, Eq. (23) becomes

$$\frac{\mathrm{d}^2 v}{\mathrm{d}\tau^2}(\tau_0) = v\frac{\mathrm{d}\pi}{\mathrm{d}\tau}(\tau_0) + (\pi - p)\frac{\mathrm{d}v}{\mathrm{d}\tau}(\tau_0).$$

Then, replacing the expression of the two terms above in equation Eq. (22) we obtain

$$\frac{\mathrm{d}^2 p}{\mathrm{d}\tau^2}(\tau_0) = \frac{1}{1+v(1+\phi_0)}\left(2\frac{\mathrm{d}\pi}{\mathrm{d}\tau}(\tau_0)(p-\pi v) + v\frac{\mathrm{d}\pi}{\mathrm{d}\tau}(\tau_0)(p-1) - \frac{2p}{v^2}\left(\frac{\mathrm{d}v}{\mathrm{d}\tau}(\tau_0)\right)^2 - \frac{p^2}{v}\frac{\mathrm{d}v}{\mathrm{d}\tau}(\tau_0)\right).$$

We use Eq. (26), and we obtain

$$\frac{\mathrm{d}^2 p}{\mathrm{d}\tau^2}(\tau_0) = -\frac{p}{v^2+v^3(1+\phi_0)}\frac{\mathrm{d}v}{\mathrm{d}\tau}(\tau_0)\left(2(p(1+v)-\pi v) + 2\frac{\mathrm{d}v}{\mathrm{d}\tau}(\tau_0) - v\right).$$

We recognize the equation Eq. (10) in the parenthesis, therefore this expression can be simplified to

$$\frac{\mathrm{d}^2 p}{\mathrm{d}\tau^2}(\tau_0) = \frac{p}{v+v^2(1+\phi_0)}\frac{\mathrm{d}v}{\mathrm{d}\tau}(\tau_0).$$

Since we assume $p < 0$ in this section, we finally have the following equivalence:

$$\frac{\mathrm{d}^2 p}{\mathrm{d}\tau^2}(\tau_0) < 0 \iff \frac{\mathrm{d}v}{\mathrm{d}\tau}(\tau_0) > 0.$$

Under the assumption $p > y_0$, the first derivative of the dimensionless volume $v$ is strictly positive. Thus we can only have a non-monotonic behavior with a local maximum (or a monotonic behavior if the first derivative does not vanish) and this is possible only if $\alpha_0 < \phi_0 y_0(\phi_0 y_0 - 1)$.

We notice that if $\alpha_0 \geq \phi_0 y_0(\phi_0 y_0 - 1)$ then the first derivative of $p$ is strictly positive when $p$ is in the interval $[p_-, 0]$. Because $p_- < y_0$, we must have $p > p_-$ in this case of growth. As a consequence, turgor pressure in the fiber increases until it reaches the value of the turgor pressure in the seed cells, thus the case $p < 0$ is no longer true and we get $p \geq 0$, which is the case investigated in next section B.2.

Repeating the analysis yields the same results for the dimensionless osmotic pressure $\pi$ – we do not present this analysis here.

***B.2. Case $P_{fiber} \geq P_{seed}$ ($p \geq 0$).*** We seek to demonstrate in this section that the dimensionless turgor pressure $p \mapsto p(\tau)$ defined over $\mathbb{R}_+$ can have at most one local minimum but no local maximum. Because we assume that $p \geq 0$, the dimensionless yield threshold $y_0$ can be either positive or negative.

We rewrite Eq. (27) using $p \geq 0$. At $\tau = \tau_0$ we have

$$\frac{\mathrm{d}p}{\mathrm{d}\tau}(\tau_0) = 0 \implies -p^2\left[1+v(1+v\phi_0)(1+\phi_0)\right] + pv\left[\phi_0 y_0 v(1+\phi_0) + (1+v\phi_0)(\phi_0 y_0 - 1)\right] + v^2(\alpha_0 + \phi_0 y_0(1-\phi_0 y_0)) = 0. \quad [33]$$

The discriminant of this last quadratic polynomial is

$$\Delta = v^2\left[\left[v\phi_0 y_0(1+\phi_0) + (1+v\phi_0)(\phi_0 y_0 - 1)\right]^2 + 4\left[\alpha_0 + \phi_0 y_0(1-\phi_0 y_0)\right]\left[1+v(1+v\phi_0)(1+\phi_0)\right]\right].$$

There are solutions to Eq. 33 only if $\Delta \geq 0$, which is equivalent to

$$v^2\phi_0\left[4\alpha_0(1+\phi_0) + \phi_0(y_0+1)^2\right] + 2v\left[2\alpha_0(1+\phi_0) + \phi_0(1+y_0)(1-y_0\phi_0)\right] + (1-\phi_0 y_0)(3\phi_0 y_0 + 1) + 4\alpha_0 \geq 0. \quad [34]$$

This is a quadratic polynomial for $v$, with discriminant $\Delta'$. If $\Delta' \leq 0$, then condition (34) is true because the coefficient of $v^2$ is strictly positive. If $\Delta' > 0$, then the polynomial in $v$ has two roots $v_1$ and $v_2$ ($v_1 < v_2$). If both $v_1$ and $v_2$ are negative, then, because we consider $v > 0$, condition (34) is true. If only one solution is positive (it would be $v_2$), then we must have $v \geq v_2$ for $Delta \geq 0$. If both solutions are positive, then we must have $v \leq v_1$ or $v \geq v_2$ for $Delta \geq 0$.

Now, we have all the conditions to have $\Delta \geq 0$. In the following, we assume the inequality to be strict as the case $\Delta = 0$ can be obtained by continuation. The roots of the polynomial in Eq. 33 are $p_-$ and $p_+$, such that

$$p_\pm = \frac{1}{2}\frac{v^2\phi_0 y_0 (1+\phi_0) + v(1+v\phi_0)(\phi_0 y_0 - 1) \pm \sqrt{\Delta}}{1 + v(1+v\phi_0)(1+\phi_0)}. \quad [35]$$

We next examine whether these roots meet the conditions of this section, *i.e.* $p(\tau_0) \geq 0$ and $p(\tau_0) > y_0$. We start with

$$p_\pm \geq 0 \iff \pm\sqrt{\Delta} \geq g(v) \text{ with } g(v) = v(1+v\phi_0)(1-\phi_0 y_0) - v^2\phi_0 y_0 (1+\phi_0). \quad [36]$$

The outcome depends on the sign of $g(v)$. If negative, then the solution $p_-$ does not meet the condition $p_- \geq 0$. For the solution $p_+$ we obtain

$$p_+(\tau_0) \geq 0 \iff \alpha_0 \geq \phi_0 y_0 (\phi_0 y_0 - 1).$$

If $g(v)$ is positive, then the solution $p_+$ satisfies the condition, and for the other solution we have

$$p_-(\tau_0) \geq 0 \iff \alpha_0 \leq \phi_0 y_0 (\phi_0 y_0 - 1).$$

Now, we consider the condition $p(\tau_0) > y_0$. We have

$$p(\tau_0) > y_0 \iff \pm\sqrt{\Delta} > f(v) \text{ with } f(v) = v^2\phi_0(1+y_0) + v(1+y_0(2+\phi_0)) + 2y_0.$$

If $f(v) > 0$, then the solution $p_-$ does not satisfy the relation $p(\tau_0) > y_0$, and for the solution $p_+$ we obtain

$$p_+(\tau_0) > y_0 \iff v^2\alpha_0 - vy_0(1+y_0) - y_0^2 > 0.$$

If $f(v) < 0$, then the solution $p_+$ satisfies the relation $p(\tau_0) > y_0$, and for the solution $p_-$ we have after calculation

$$p_-(\tau_0) > y_0 \iff v^2\alpha_0 - vy_0(1+y_0) - y_0^2 < 0.$$

We remark that

$$f(v) < 0 \implies y_0 < 0 \implies g(v) < 0.$$

Thus, we can not have simultaneously for the same dimensionless volume $v$ the condition $f(v) < 0$ and $g(v) > 0$, thus the solution $p_-$ does not meet simultaneously the conditions $p_-(\tau_0) > y_0$ and $p_-(\tau_0) > 0$. The only solution for the case $\Delta > 0$ to have the first derivative equal to zero is therefore $p_+$, under some conditions on $v$ according to the value of the parameters.

In what follows we study the sign of the second derivative of $p$. Taking into account that $p \geq 0$, the second derivative of the dimensionless osmotic pressure $\pi$ (Eq. 24) can be rewritten as

$$\frac{\mathrm{d}^2\pi}{\mathrm{d}\tau^2} = -\left(\pi - p\right)\frac{\mathrm{d}\pi}{\mathrm{d}\tau} - \left(\pi + 1\right)\left(\frac{\mathrm{d}\pi}{\mathrm{d}\tau} - \frac{\mathrm{d}p}{\mathrm{d}\tau}\right).$$

The second derivative of the dimensionless volume $v$ (Eq. 23) becomes

$$\frac{\mathrm{d}^2 v}{\mathrm{d}\tau^2} = v\frac{\mathrm{d}\pi}{\mathrm{d}\tau} + \left(\pi - p\right)\frac{\mathrm{d}v}{\mathrm{d}\tau} - \left(v + 1\right)\frac{\mathrm{d}p}{\mathrm{d}\tau}.$$

At $\tau = \tau_0$, these equations become

$$\frac{\mathrm{d}^2\pi}{\mathrm{d}\tau^2}\left(\tau_0\right) = \frac{\mathrm{d}\pi}{\mathrm{d}\tau}\left(\tau_0\right)\left(-2\pi - 1 + p\right)$$

and

$$\frac{\mathrm{d}^2 v}{\mathrm{d}\tau^2}\left(\tau_0\right) = v\frac{\mathrm{d}\pi}{\mathrm{d}\tau}\left(\tau_0\right) + \left(\pi - p\right)\frac{\mathrm{d}v}{\mathrm{d}\tau}\left(\tau_0\right).$$

We can calculate the second derivative of $p$ (Eq. 22), and compare its value to zero

$$\frac{\mathrm{d}^2 p}{\mathrm{d}\tau^2}\left(\tau_0\right) < 0 \iff p\frac{\mathrm{d}v}{\mathrm{d}\tau}\left(\tau_0\right)\left[p\left(2 + v\left(1 + \phi_0\right)\right) + v\left(1 - \phi_0 y_0\right)\right] < 0.$$

The case $p = 0$ does not meet this condition. Because we assumed that the fiber is growing, $\mathrm{d}v/\mathrm{d}\tau(\tau_0) > 0$, thus the inequality above implies that

$$p\left(\tau_0\right) < \frac{v\left(\tau_0\right)\left(\phi_0 y_0 - 1\right)}{2 + v\left(\tau_0\right)\left(1 + \phi_0\right)}. \quad [37]$$

Because we assumed that $p$ is positive, the quantity $\phi_0 y_0 - 1$ must be positive, we therefore have $\phi_0 y_0 > 1$ which also implies that $y_0 > 0$ because $\phi_0 > 0$. The solution $p_+$ must verify Eq. (37) for the second derivative of $p$ to be negative, which is equivalent to

$$\sqrt{\Delta} < q(v) \text{ with } q(v) = -v^3\phi_0\left(1 + \phi_0\right)\left(1 + y_0\right) - v^2\left(\phi_0\left(y_0\left(1 + 3\phi_0\right) - 1\right) + 1\right).$$

If $q\left(v\right) < 0$, the $p_+\left(\tau_0\right)$ does not verify inequality (37). If $q\left(v\right) > 0$ then we obtain

$$\begin{aligned}\sqrt{\Delta} < q\left(v\right) \iff & v^2\alpha_0\left(1 + \phi_0\right)^2 + 2v\left[2\alpha_0\left(1 + \phi_0\right) + \phi_0\left(1 - \phi_0 y_0\right)\left(1 + y_0\right)\right] \\ & + 4\alpha_0 + \left(3\phi_0 y_0 + 1\right)\left(1 - \phi_0 y_0\right) < 0.\end{aligned}$$

We assumed that $\phi_0 y_0 > 1$ thus $y_0 > 0$. We thus find that

$$q\left(v\right) > 0 \iff v < \frac{-\left(\phi_0\left(y_0\left(1 + 3\phi_0\right) - 1\right) + 1\right)}{\phi_0\left(1 + \phi_0\right)\left(1 + y_0\right)}.$$

Since $y_0 > 0$ and $v > 0$ we have the following implication

$$q\left(v\right) > 0 \implies \phi_0 y_0 < \frac{\phi_0 - 1}{1 + 3\phi_0}.$$

But we assumed that $\phi_0 y_0 > 1$ for (37), and

$$\phi_0 y_0 > 1 \implies \phi_0 y_0 > \frac{\phi_0 - 1}{1 + 3\phi_0}.$$

Thus we obtain a contradiction, and the case $q\left(v\right) > 0$ is not compatible with Eq. (37). Thus the solution $p_+$ does not verify the condition (37), and the second derivative of $p$ can not be negative.

A similar analysis shows that it is possible to have the second derivative of $p$ positive with the solution $p_+$.

**B.3. Relations with the initial conditions.** Here we consider whether the initial conditions enable an extremum of pressure. In the preceding section, we investigated the sign of the second derivative of $p$. We obtained the value of $p$ when the first derivative is equal to zero (Eq. 35). We can also calculate the isocline of $p$ in the phase space $(p, v)$ and show that this isocline is strictly monotonic, increasing for $p \geq 0$ and decreasing for $p < 0$. This leads to a necessary condition for the first derivative to vanish at a certain time $\tau_0$. Indeed, for $p$ to have a local maximum the initial condition $p(0)$ must be lower than the maximum reached, and for the local minimum, the initial condition must be higher than the minimum reached (i.e. when the first derivative equals to zero). We note that we also obtained the same results for $\pi$.

In the case $p < 0$, in order to have the second derivative of the turgor pressure negative, it is necessary to have $p\,(0) < p_{\max}$ such that

$$p_{\max} = \frac{1}{2v^2}\frac{v\phi_0 y_0\,(v + v\phi_0 - 1) + v\,(1 + v\phi_0)\,(\phi_0 y_0 - 1) + \sqrt{\Delta}}{\phi_0\,(1+\phi_0)}, \qquad [38]$$

with $\Delta$ defined as

$$\Delta = v^2\left[\left[\phi_0 y_0\,(v + v\phi_0 - 1) + (1 + v\phi_0)\,(\phi_0 y_0 - 1)\right]^2 + 4v^2\left[\alpha_0 + \phi_0 y_0\,(1 - \phi_0 y_0)\right]\left[\phi_0\,(1+\phi_0)\right]\right].$$

In the case $p \geq 0$, in order to have the second derivative of the turgor pressure positive, it is necessary to have $p\,(0) > p_{\min}$ such that

$$p_{\min} = \frac{1}{2}\frac{v^2\phi_0 y_0\,(1+\phi_0) + v\,(1 + v\phi_0)\,(\phi_0 y_0 - 1) + \sqrt{\Delta}}{1 + v\,(1 + v_0\phi_0)\,(1+\phi_0)}, \qquad [39]$$

with $\Delta$ defined as

$$\Delta = v^2\left[\left[v\phi_0 y_0\,(1+\phi_0) + (1 + v\phi_0)\,(\phi_0 y_0 - 1)\right]^2 + 4\left[\alpha_0 + \phi_0 y_0\,(1 - \phi_0 y_0)\right]\left[1 + v\,(1 + v\phi_0)\,(1+\phi_0)\right]\right].$$

For the same reasons, in order to have the second derivative of the osmotic pressure negative, for the case $p < 0$, it is necessary to have $\pi\,(0) < \pi_{\max}$ such that

$$\pi_{\max} = \frac{\phi_0\,(y_0\,(1+v) - v) - 1 + \sqrt{\Delta}}{2v\phi_0}, \qquad [40]$$

with $\Delta$ defined as

$$\Delta = \left[\phi_0\,(y_0\,(1+v) - v) - 1\right]^2 + 4v\phi_0\left[\alpha_0\,(1 + v\,(1+\phi_0)) + v\phi_0 y_0\right].$$

Finally, in order to have the second derivative of the osmotic pressure positive, for the case $p \geq 0$, it is necessary to have $\pi\,(0) > \pi_{\min}$ such that

$$\pi_{\min} = \frac{v\phi_0\,(y_0 - 1) - 1 + \sqrt{\Delta}}{2\,(v\phi_0 + 1)}, \qquad [41]$$

with $\Delta$ defined as

$$\Delta = 4\alpha_0\,(\phi_0 v + 1)\,(v\,(\phi_0 + 1) + 1) + (v\phi_0\,(y_0 + 1) + 1)^2.$$

Moreover, because the osmotic pressure and the turgor pressure must be positive to have a physical meaning, we must have $\pi \geq -1$, which implies $p \geq \frac{-v}{v+1}$ if $p > y_0$ or $p \geq \frac{v(\phi_0 y_0 - 1)}{v(1+\phi_0)+1}$ if $p \leq y_0$ and $P_{\text{fiber}} > 0$ implies $p \geq -\frac{P_{\text{seed}}}{\pi_{\text{seed}}}$.

This naturally adds conditions on the parameters. For instance, for the minimum possible value of $\pi$, i.e. $\pi = -1$, to have the system has the osmotic pressure and the turgor pressure positive, we must have:

$$\phi_0 \geq \frac{1 - \frac{P_{\text{seed}}}{\pi_{\text{seed}}}}{y_0 + \frac{P_{\text{seed}}}{\pi_{\text{seed}}}}$$

or in other terms:

$$\phi_0 \geq \frac{\pi_{\text{seed}} - P_{\text{seed}}}{Y}.$$

We can deduce such relations between the parameters by considering the limit of $p$ and $\pi$ at $+\infty$. These relations have to be verified if we want to have the turgor pressure and osmotic pressure positive in numerical simulations.

At this point, we have analysed all possible behaviors in terms of temporal variations of the three dimensionless variables $v$, $\pi$, and $p$. These behaviors and the corresponding conditions are summarised in the following section.

#### 3. Summary of the analytical study and numerical illustrations

The different possible behaviors for the pressures and the volume with the model are detailed in the following figures and tables.

**Table S3. Behaviors of the model. the quantities $p_{\max}$, $p_{\min}$, $\pi_{\max}$, $\pi_{\min}$ are defined previously in equations (38,39,40,41) and depend on the initial dimensionless volume $v(0)$ and on the three dimensionless parameters $\alpha_0$, $\phi_0$ and $y_0$. The variables $v$, $p$ and $\pi$ represent respectively the dimensionless volume, turgor pressure and osmotic pressure.**

| | No growth $p(\tau) \le y_0$ | Growth $p(\tau) > y_0$ | | | | | |
|---|---|---|---|---|---|---|---|
| | | 1 - Low yield threshold and low osmotic adjustment $y_0 < 0$ and $\alpha_0 < \phi_0 y_0 (\phi_0 y_0 - 1)$ | | | | 2 - High yield threshold or High osmotic adjustment $y_0 \ge 0$ or $\alpha_0 \ge \phi_0 y_0 (\phi_0 y_0 - 1)$ | |
| $v$ | Constant | Monotonic (strict growth) | | | | Monotonic (strict growth) | |
| $p$ | monotonic (strict growth or strict degrowth) | A - Low initial turgor pressure $p(0) < p_{\max}$ | | B - High initial turgor pressure $p(0) \ge p_{\max}$ | | C - Low initial turgor pressure $p(0) \le p_{\min}$ | D- High initial turgor pressure $p(0) > p_{\min}$ |
| | | Non-monotonic (local maximum) or monotonic (strict growth) | | Monotonic (strict degrowth) | | Monotonic (strict growth) | Non-monotonic (local minimum) or monotonic (strict degrowth) |
| $\pi$ | Monotonic (strict growth or strict degrowth) | 3 - Low osmotic adjustment $\alpha_0 < \frac{1-y_0\phi_0}{\phi_0}$ | | 4 - High osmotic adjustment $\alpha_0 \ge \frac{1-y_0\phi_0}{\phi_0}$ | | C - Low initial osmotic pressure $\pi(0) \le \pi_{\min}$ | D - High initial osmotic pressure $\pi(0) > \pi_{\min}$ |
| | | A - Low initial osmotic pressure $\pi(0) < \pi_{\max}$ | B - High initial osmotic pressure $\pi(0) \ge \pi_{\max}$ | A - Low initial osmotic pressure $\pi(0) < \pi_{\max}$ | B - High initial osmotic pressure $\pi(0) \ge \pi_{\max}$ | | |
| | | Non-monotonic (local maximum) or monotonic (strict growth) | Monotonic (strict degrowth) | monotonic (strict growth) | Non-monotonic (local minimum) or monotonic (strict degrowth) | Monotonic (strict growth) | Non-monotonic (local minimum) or monotonic (strict degrowth) |

**Table S4. Values used for the curves in Figure S2. All the cases from Table S3 are illustrated. In all cases, we fixed the following parameters: $L_r = 0.1\, MPa^{-1}.h^{-1}$, $\mu = 0.001\, mm^3 MPa^{-1}.h^{-1}$, $\pi_{\mathbf{seed}} = 1\, MPa$, $P_{\mathbf{seed}} = 0.18\, MPa$, $Y = 0.06\, MPa$, and $\phi = 1.2\, MPa^{-1}.h^{-1}$. Therefore $\phi_0 y_0 (\phi_0 y_0 - 1) = 3.51$ and $1 - y_0\phi_0/\phi_0 = 0.20$; we state in this Table whether these quantities are higher or lower than $\alpha_0$. In the fist column, $P_{\mathbf{fiber}}$ or nd $\pi_{\mathbf{fiber}}$ indicates that, respectively, turgor or osmotic pressures are shown. $\alpha$ is given in units of $MPa.h^{-1}$, while the initial condition $P(0)$ is in $MPa$.**

| Condition | $\alpha$ | $P(0)$ | $\alpha_0$ | $\phi_0 y_0 (\phi_0 y_0 - 1)$ | $\frac{1-y_0\phi_0}{\phi_0}$ |
|---|---|---|---|---|---|
| 1-A-$P_{\text{fiber}}$ (figure S2a) | $0.3$ | $0.16$ | $3.0$ | Higher | Lower |
| 1-B-$P_{\text{fiber}}$ (figure S2b) | $0.3$ | $0.18$ | $3.0$ | Higher | Lower |
| 1-3-A-$\pi_{\text{fiber}}$ (figure S2c) | 0.01 | 0.155 | 0.1 | Higher | Higher |
| 1-3-B-$\pi_{\text{fiber}}$ (figure S2d) | 0.01 | 0.16 | 0.1 | Higher | Higher |
| 1-4-A-$\pi_{\text{fiber}}$ (figure S2e) | 0.3 | 0.155 | 3.0 | Higher | Lower |
| 1-4-B-$\pi_{\text{fiber}}$ (figure S2f) | 0.3 | 0.18 | 3.0 | Higher | Lower |
| 2-C-$P_{\text{fiber}}$ (figure S2g) | 0.4 | 0.16 | 4.0 | Lower | Lower |
| 2-D-$P_{\text{fiber}}$ (figure S2h) | 0.4 | 0.20 | 4.0 | Lower | Lower |
| 2-C-$\pi_{\text{fiber}}$ (figure S2i) | 0.5 | 0.16 | 4.0 | Lower | Lower |
| 2-D-$\pi_{\text{fiber}}$ (figure S2j) | 0.5 | 0.19 | 5.0 | Lower | Lower |

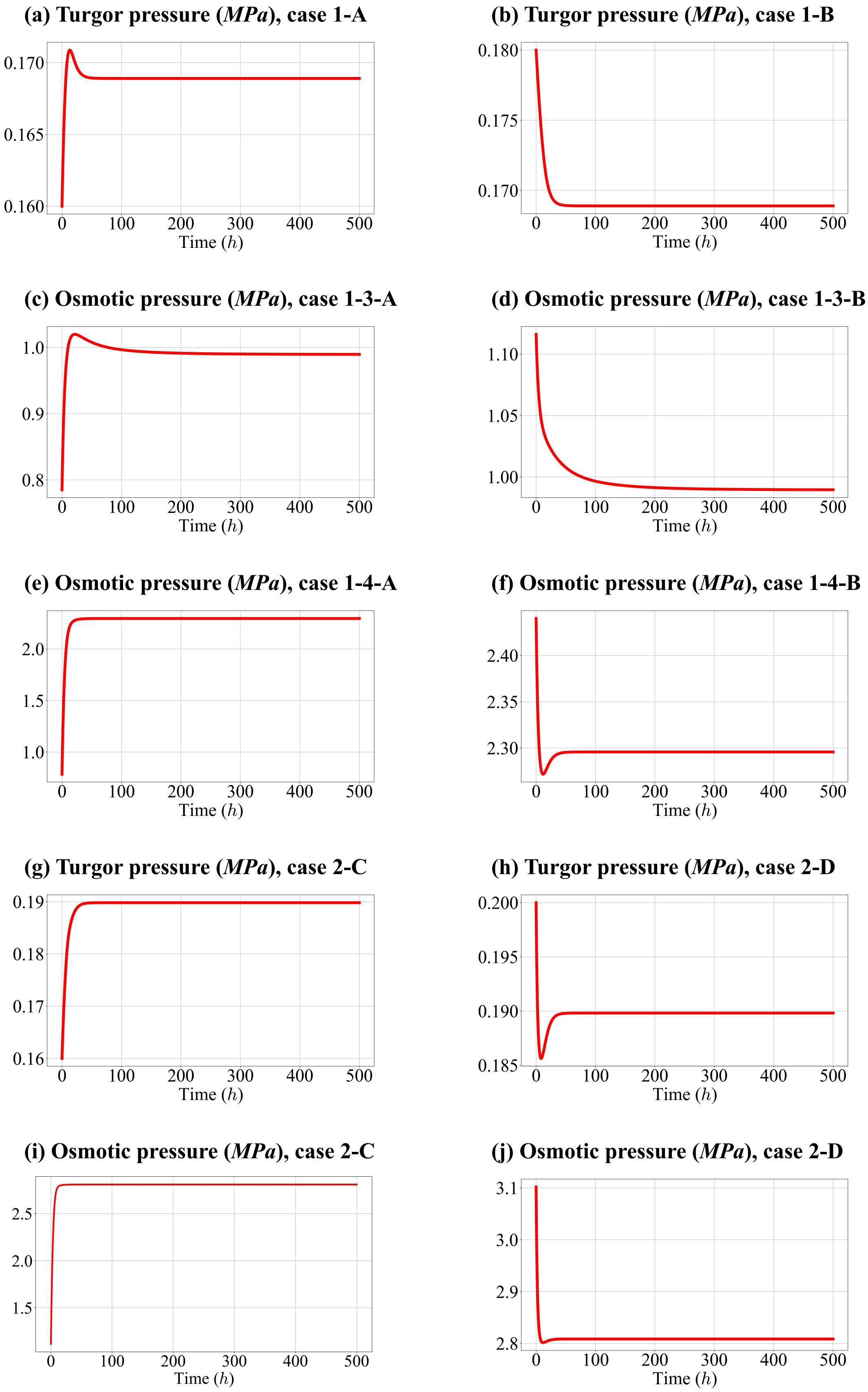

**Fig. S2. Behaviors of the osmotic and turgor pressure according to the cases reported in Table S3. The values of the parameters used are given in Table S4.**

#### 4. Condition for a peak of turgor pressure and biologically-relevant parameters

We demonstrated that, in order to have a peak of turgor pressure, we need to have $\alpha_0 < \phi_0 y_0 (\phi_0 y_0 - 1)$ and $y_0 < 0$. In figure S3 we plot this condition in the dimensionless parameter space using parameter values reported in table S2. We note that the values of $\phi_0 y_0 (\phi_0 y_0 - 1)$ are very low compared to the biophysical value reported for $\alpha_0$. The necessary conditions $\alpha_0 < \phi_0 y_0 (\phi_0 y_0 - 1)$ and $y_0 < 0$ can be satisfied if the value of $\alpha_0$ is lower than approximately $6.0 \times 10^{-4}$, which represents 1.7% of the biophysical range reported of $\alpha_0$.

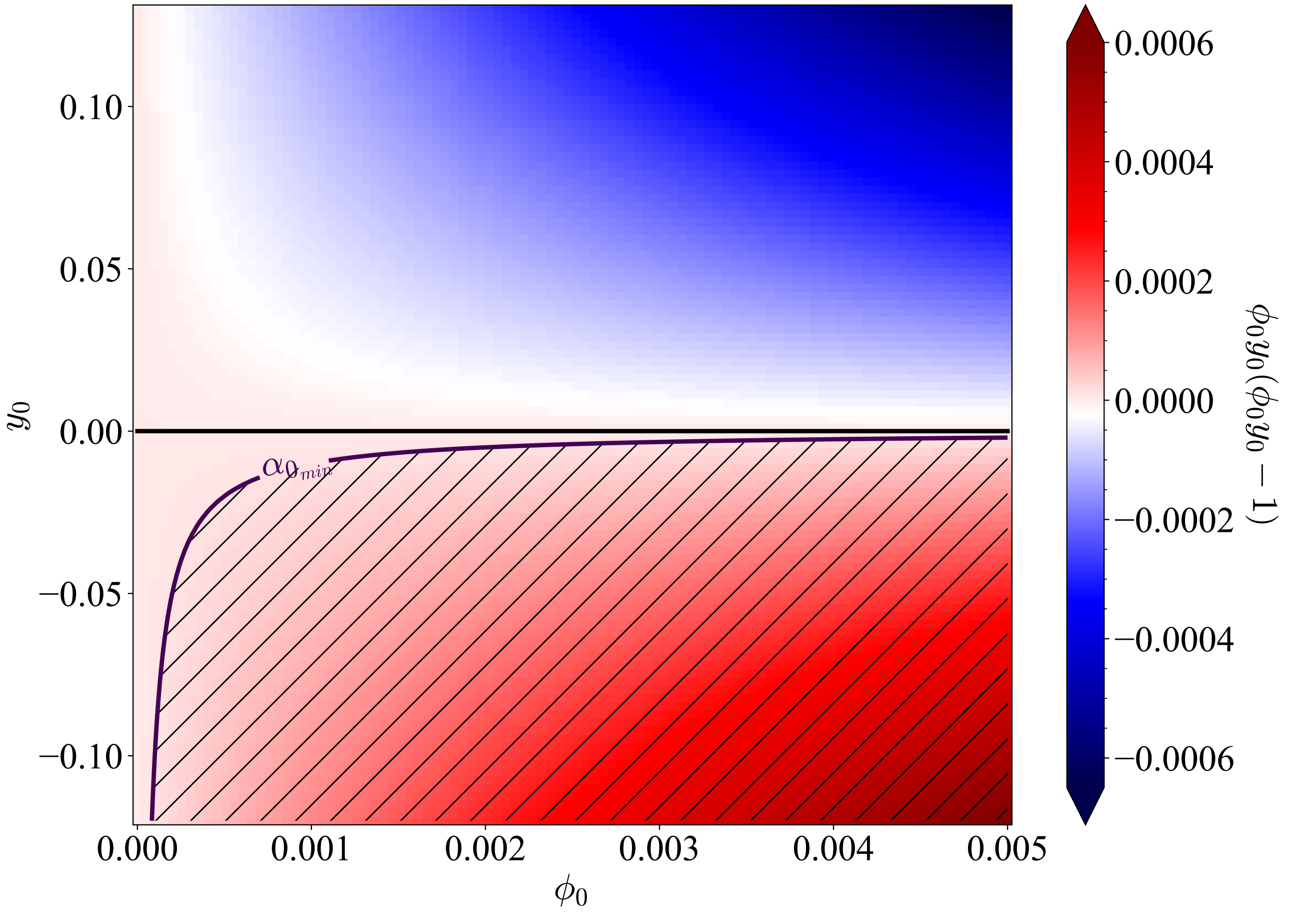

**Fig. S3. Condition $\alpha_0 \geq \phi_0 y_0 (\phi_0 y_0 - 1)$ in the space of dimensionless parameters $\phi_0$ and $y_0$. The hatched area corresponds to the region of the parameter space where the condition is satisfied, for $\alpha_{0_{\mathrm{min}}} = 1.0 \times 10^{-5}$ as computed from Table S1, and the continuous purple line corresponds to this value of $\alpha = \alpha_{0_{\mathrm{min}}}$.**

#### 5. Details of the sensitivity analysis

As part of the sensitivity analysis of the model near reference parameters, we show in Figures S4-S6 the ratio between the final value of the observable variable and its value for the reference parameters.

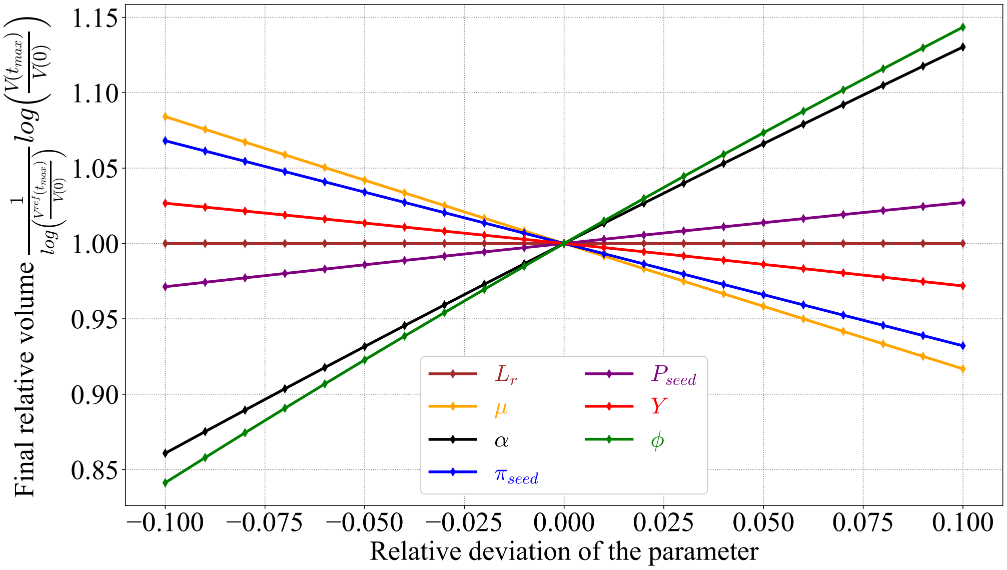

**Fig. S4. Sensitivity of the volume to the seven parameters of the model. Normalised value plotted as function of the relative change of the parameter.**

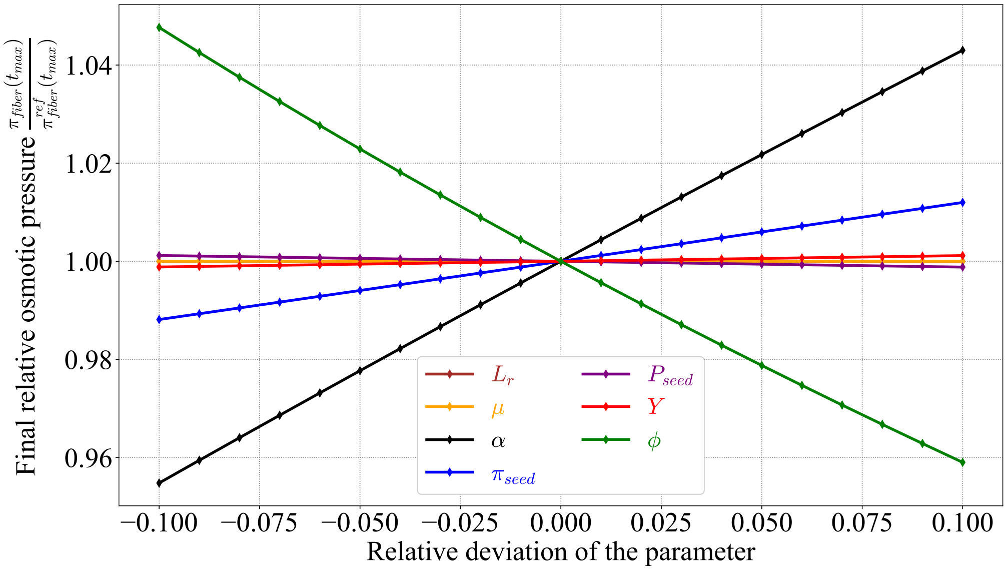

**Fig. S5. Sensitivity of the osmotic pressure to the seven parameters of the model. Normalised value plotted as function of the relative change of the parameter.**

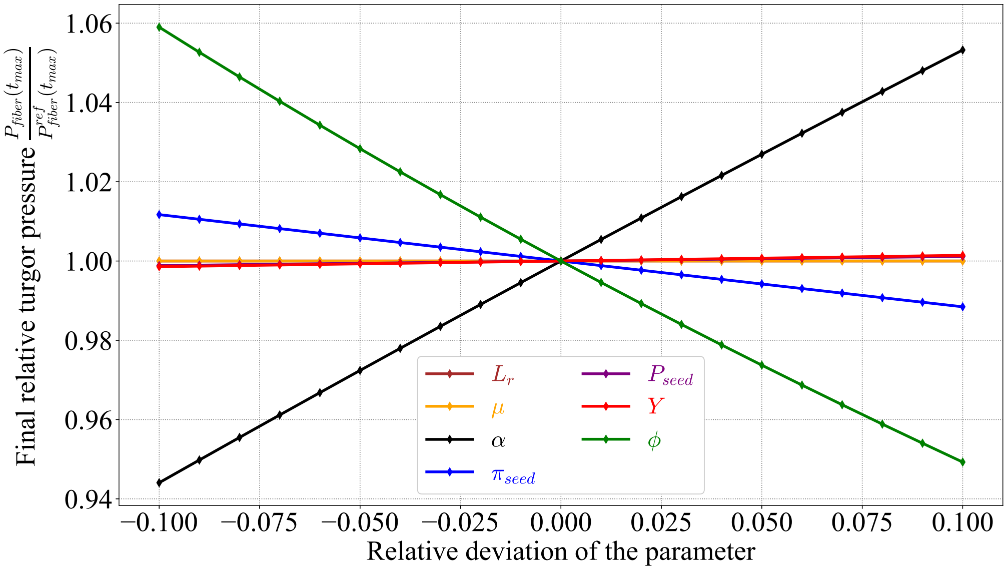

**Fig. S6. Sensitivity of the turgor pressure to the seven parameters of the model. Normalised value plotted as function of the relative change of the parameter.**

#### 6. Dynamic source of solute

In Figure S7 we plot the behavior of the volume and pressures with a dynamic source of solute $\alpha$ corresponding to the temporal pattern shown in Panel a.

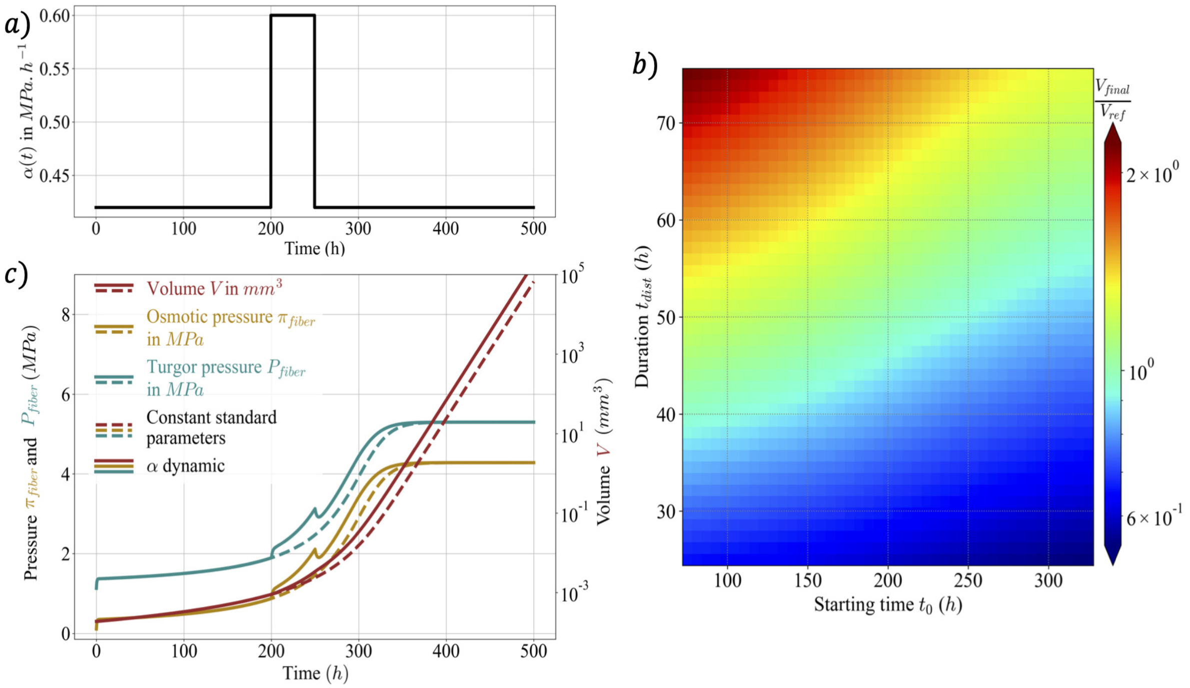

**Fig. S7. Pressures and volume with a dynamic source of solute $\alpha$. a) Source of solute $\alpha$ as a function of time. b) Normalised final volume as function of the start time $t_0$ and of the duration $t_{dist}$ of the period during which the source is enhanced (the normalised volume corresponds to the ratio of the final volume at $3000\ h$ to the final volume at the same time taking $t_{dist} = 50\ h$ and $t_0 = 200\ h$)). c) Continuous lines correspond to the behavior of the model with dynamic source. Dashed lines correspond to the case when all parameters are constant and equal to their reference values.**
